# CEP78 functions downstream of CEP350 to control biogenesis of primary cilia by negatively regulating CP110 levels

**DOI:** 10.1101/2020.10.05.325936

**Authors:** André Brás Gonçalves, Sarah Kirstine Hasselbalch, Beinta Biskopstø Joensen, Sebastian Patzke, Pernille Martens, Signe Krogh Ohlsen, Mathieu Quinodoz, Konstantinos Nikopoulos, Reem Suleiman, Magnus Per Damsø Jeppesen, Catja Weiss, Søren Tvorup Christensen, Carlo Rivolta, Jens S. Andersen, Pietro Farinelli, Lotte Bang Pedersen

**Affiliations:** Department of Biology, Section for Cell Biology and Physiology, University of Copenhagen, Universitetsparken 13, DK-2100 Copenhagen Ø, Denmark; Department of Radiation Biology, Institute for Cancer Research, Norwegian Radium Hospital, Oslo University Hospital, Oslo, Norway; Department of Biochemistry and Molecular Biology, University of Southern Denmark, Campusvej 55, 5230 Odense M, Denmark; Institute of Molecular and Clinical Ophthalmology Basel (IOB), Basel, Switzerland; Department of Ophthalmology, University of Basel, Basel, Switzerland; Department of Genetics and Genome Biology, University of Leicester, Leicester, United Kingdom; Department of Computational Biology, University of Lausanne, Switzerland; Instituto de Medicina Molecular João Lobo Antunes, Faculdade de Medicina, Universidade de Lisboa, Lisbon, Portugal

## Abstract

CEP78 is a centrosomal protein implicated in ciliogenesis and ciliary length control, and mutations in the *CEP78* gene cause retinal cone-rod dystrophy associated with hearing loss. However, the mechanism by which CEP78 affects cilia formation is unknown. Based on a recently discovered disease-causing *CEP78* p.L150S mutation, we identified the disease-relevant interactome of CEP78. We confirmed that CEP78 interacts with the EDD1-DYRK2-DDB1^VPRBP^ E3 ubiquitin ligase complex, which is involved in CP110 ubiquitination and degradation, and identified a novel interaction between CEP78 and CEP350 that is weakened by the CEP78^L150S^ mutation. We show that CEP350 promotes centrosomal recruitment and stability of CEP78, which in turn leads to centrosomal recruitment of EDD1. Consistently, cells lacking CEP78 display significantly increased cellular and centrosomal levels of CP110, and depletion of CP110 in CEP78-deficient cells restored ciliation frequency to normal. We propose that CEP78 functions downstream of CEP350 to promote ciliogenesis by negatively regulating CP110 levels via an EDD1-dependent mechanism.

## Introduction

Primary cilia are antenna-like sensory organelles that play pivotal roles in coordinating various signaling pathways important for human development and tissue homeostasis [1]. They comprise a microtubule-based axoneme core, which extends directly from the mother centriole-derived basal body and is surrounded by a bilayer membrane enriched for specific receptors, ion channels and lipids that endow the organelle with unique signaling properties [2]. Not surprisingly, mutations in genes that affect assembly, structure or function of cilia are causative for a growing number of pleiotropic diseases and syndromes called ciliopathies, which include cone-rod dystrophy in the retina and hearing loss (CRDHL; MIM# 617236) amongst others [3]. Ciliopathy genes include those coding for components of the centrosome, which contains the daughter and mother centriole and gives rise to the ciliary basal body. The mother centriole is distinguished from the daughter centriole by distal and subdistal appendages, which play critical roles in vesicle docking at the onset of ciliogenesis and in microtubule anchoring, respectively. In addition, the centrosome contains pericentriolar material and is associated with centriolar satellites that affect cilia biogenesis and function in various ways [4].

Assembly of primary cilia is a complex, multistep process that is tightly coordinated with the cell cycle. In actively-proliferating cells, centriolar coiled coil protein 110 (CP110; also known as CCP110) and centrosomal protein of 97 kDa (CEP97) cap the distal ends of both mother and daughter centrioles and suppress ciliogenesis. Furthermore, overexpression of CP110 in growth-arrested cells prevents ciliogenesis [5]. CP110 also regulates centriole duplication and length control during S phase and interacts with key regulators of centriole duplication, including PLK4 [6–9]. As cells enter G1/G0, ciliogenesis begins with recruitment of pre-ciliary vesicles to the distal end of the mother centriole. The vesicles subsequently fuse to form a larger ciliary vesicle underneath which the ciliary transition zone and axoneme are assembled. The axoneme is further extended by intraflagellar transport (IFT), and the ciliary vesicle expands and matures to form the ciliary membrane and a surrounding sheath that fuses with the plasma membrane upon completion of ciliogenesis [10–12].

Initiation of transition zone formation and axoneme extension during early stages of ciliogenesis require removal of the CEP97-CP110 complex from the distal end of the mother centriole [5]. This process relies on M-Phase Phosphoprotein 9 (MPP9), which interacts directly with CEP97 and cooperates with kinesin KIF24 to recruit the CEP97-CP110 complex to the distal centriole end [13, 14]. During ciliogenesis, phosphorylation of MPP9 by Tau tubulin kinase 2 (TTBK2) leads to degradation of MPP9 by the ubiquitin proteasome system (UPS), which results in destabilization of the CEP97-CP110 complex causing its removal from the distal end of the mother centriole [14]. TTBK2 is recruited to the mother centriole distal appendages by CEP164 [15, 16] where it also phosphorylates CEP83 to promote CP110 removal [17]. Consequently, depletion of CEP164, TTBK2, CEP83 or other centriole distal appendage proteins that regulate their localization and/or function impairs ciliogenesis [18–23]. Mother centriole recruitment of TTBK2 and removal of the distal CEP97-CP110 cap additionally requires the subdistal CEP350-FOP-CEP19 complex, which furthermore interacts with RABL2 and IFT-B complex components to promote their axonemal entry [24–26].

Despite recent advances, the precise mechanisms by which CP110 regulates ciliogenesis and is removed from the mother centriole at the onset of ciliogenesis remain incompletely understood. For example, a recent study implicated the homologous to the E6AP carboxyl terminus (HECT) type EDD1-DYRK2-DDB1^VPRBP^ E3 ligase complex in ubiquitination and degradation of CP110 via a mechanism involving direct interaction of viral protein R binding protein (VPRBP; also known as DCAF1) and centrosomal protein of 78 kDa (CEP78) [27]. Specifically, the authors reported that CEP78 suppresses CP110 ubiquitination by EDD1 (also known as UBR5 and EDD) and it was proposed that CP110 is phosphorylated by DYRK2 and thereby recognized and brought close to EDD1 by VPRBP. EDD1 then transfers ubiquitin to CP110 leading to its degradation. When CEP78 binds VPRBP it induces a conformational change in the complex thereby preventing CP110 ubiquitination. Further, they demonstrated that depletion of CEP78 promoted centriole elongation, whereas *CEP78* over-expression inhibited primary cilia formation in hTERT-immortalized retinal pigment epithelial (RPE1) cells [27]. On the other hand, knockout (KO) of *Dyrk2* in the mouse was reported to result in elongation of primary cilia, but centrosomal CP110 levels appeared unaffected in *Dyrk2* mouse KO cells [28]. Therefore, it remains uncertain how CEP78 and the EDD1-DYRK2-DDB1^VPRBP^ complex affect CP110 homeostasis to control ciliogenesis.

*CEP78* is composed of 16 exons and encodes a protein comprising 722 amino acids that possesses five consecutive leucine rich repeats (LRR) at the N-terminus, and a coiled-coil (CC) domain at the C-terminus [29, 30]. Four independent studies have reported eight different mutations in *CEP78* in patients with CRDHL [29–32], whereas one study identified a homozygous *CEP78*-truncating variant in a family with non-syndromic retinitis pigmentosa (MIM# 268003; [33]), another form of retinal degeneration. Of these studies, two have investigated the functional consequences of human *CEP78* mutations at the cellular level. In one study, whole-exome sequencing (WES) identified biallelic mutations in *CEP78* in two unrelated families from Greece and Sweden, respectively [29]. The Greek subject had a homozygous base substitution at the splice donor site in intron 3 of *CEP78* (NM_032171.2:c.499+1G>T). Two subjects from a Swedish family carried heterozygous mutations, one base substitution in intron 3 (NM_032171.2:c.499+5G>A) and a single nucleotide deletion in exon 5 (NM_032171.2:c.633del; p.Trp212GlyfsTer18) causing a frameshift. These *CEP78* mutations lead to exon skipping and premature stop codons accompanied by almost undetectable levels of CEP78 protein in human skin fibroblasts (HSFs) of affected individuals. Furthermore, it was found that HSFs from these patients have significantly longer primary cilia as compared to control cells [29]. More recently, a missense mutation in *CEP78* (NM_032171.2:c.449T>C; p.Leu150Ser) was identified in three unrelated families from Belgium and Germany diagnosed with CRDHL [30]. In the two Belgian families, affected individuals were homozygous for the p.Leu150Ser variant, whereas affected individuals from the German family displayed compound heterozygosity for this variant and a novel splice site variant, NM_032171.2:c.1462-1G>T. In all cases, HSFs from patients harboring the p.Leu150Ser mutation (hereafter: L150S) displayed decreased cellular levels of CEP78 and significantly elongated cilia compared to control HSFs [30], as seen in patient-derived HSFs with CEP78 truncating mutations [29]. In addition, other ciliopathy features were reported in some of the affected individuals with the CEP78^L150S^ mutation, including obesity, respiratory problems, diabetes 2 and infertility [30]. In summary, available data derived from human patients indicates that depletion of CEP78 leads to elongation of primary cilia at the cellular level, which manifests in ciliopathy phenotypes at the organism level. However, the mechanism by which CEP78 regulates ciliary length is not known.

In addition to the patient studies described above, some studies have addressed CEP78 function in different cell culture models. First, a study showed that siRNA-mediated depletion of CEP78 in RPE1 cells reduces the frequency of cells forming primary cilia, possibly due to centriolar anchoring defects, but the length of the residual cilia was not assessed. Similarly, depletion of CEP78 in *Planarians* was shown to impair motile cilia formation due to defective docking of centrioles to the cell surface [34]. Another study identified CEP78 interaction with PLK4 and implicated CEP78 in PLK4-mediated centriole over duplication, whereas possible roles for CEP78 in relation to cilia were not addressed [35]. Finally, as mentioned above, a study reported that CEP78 directly interacts with VPRBP of the EDD1-DYRK2-DDB1^VPRBP^ E3 ligase complex to suppress CP110 ubiquitination and thereby stabilize it [27]. How such CEP78-mediated stabilization of CP110 might lead to the long cilia phenotype observed in patient fibroblasts lacking CEP78 [29, 30] and/or the reduced ciliation frequencies seen in CEP78-depleted *Planarians* and RPE1 cells [34] is unclear.

Here were show that in RPE1 cells and patient-derived HSFs loss of CEP78 leads to reduced ciliation frequency as well as increased length of the cilia that do form. Further, we find that a mutant line expressing a partially functional, truncated version of CEP78 displays reduced ciliation frequency but normal length of residual cilia. By taking advantage of the recently identified disease-causing CEP78^L150S^ mutation [30], we used a quantitative mass spectrometry-based approach to identify a disease-relevant interactome of CEP78. We confirmed that CEP78 interacts with the EDD1-DYRK2-DDB1^VPRBP^ complex implicated in CP110 ubiquitination and degradation [27], and furthermore identified a novel interaction between CEP78 and the N-terminal region of CEP350. The interaction of CEP78 with both VPRBP and CEP350, as well as centrosomal recruitment of CEP78, are dramatically reduced by the CEP78^L150S^ mutation. Lack of centrosomal recruitment of CEP78^L150S^ is likely due to impaired interaction with CEP350 since centrosomal and cellular levels of CEP78 were significantly decreased in *CEP350* KO cells. Conversely, cells lacking CEP78 displayed significantly decreased centrosomal levels of EDD1 and unaltered or slightly increased centrosomal levels of VPRBP and CEP350. In addition, CEP78-deficient cells showed significantly increased cellular and centrosomal levels of CP110, presumably owing to the reduced centrosomal levels of EDD1 observed in these cells. Depletion of CP110 in CEP78-deficient cells restored ciliation frequency, but not the increased length of remaining cilia, to normal. Collectively our results suggest that CEP78 functions downstream of CEP350 to promote cilia biogenesis by negatively regulating CP110 levels via an EDD1-dependent mechanism, but suppresses ciliary elongation independently of CP110.

## Results

### Depletion of CEP78 leads to fewer and longer primary cilia in cultured cells

Previous studies showed that patient-derived fibroblasts lacking CEP78 display significantly elongated primary cilia compared to control cells [29, 30] whereas depletion of CEP78 in *Planarians* and RPE1 cells was shown to significantly reduce cilia numbers [34]. To reconcile these seemingly contradictory findings we analyzed cilia numbers and length in serum-deprived wild type (WT) RPE1 cells and four different *CEP78* KO clones generated by CRISPR/Cas9 methodology. We first confirmed by western blot analysis with CEP78-specific antibody that endogenous CEP78 is lacking in three of the mutant clones, designated #2, #52 and #73, whereas clone #44 expresses a shorter version of CEP78 (Figure 1-figure supplement 1A). Sequencing of clone #44 indicated that it contains an insertion of the px459-Cas9 plasmid in exon 1 of *CEP78*, suggestion that a shorter version of CEP78 lacking the extreme N-terminus is expressed in this strain, possibly from an alternative promoter. Immunofluorescence microscopy (IFM) analysis with antibodies against ciliary (ARL13B) and centrosomal (CEP350) markers showed that the frequency of ciliated cells is significantly reduced in all four *CEP78* KO clones compared to WT cells (Figure 1A, B; Figure 1-Figure supplement 1B). In addition, the length of the remaining cilia was significantly increased in the three *CEP78* KO clones that completely lack CEP78, but identical to the average WT cilia size in clone #44 (Figure 1A, C; Figure 1-figure supplement 1C). The latter result indicates that the truncated CEP78 protein expressed in clone #44 functions normally with respect to ciliary length control, in turn suggesting that CEP78 may affect ciliogenesis and ciliary length by distinct mechanisms. For the rest of this manuscript RPE1 *CEP78* KO cells refer to clone #73 unless otherwise indicated.

**Figure 1.**
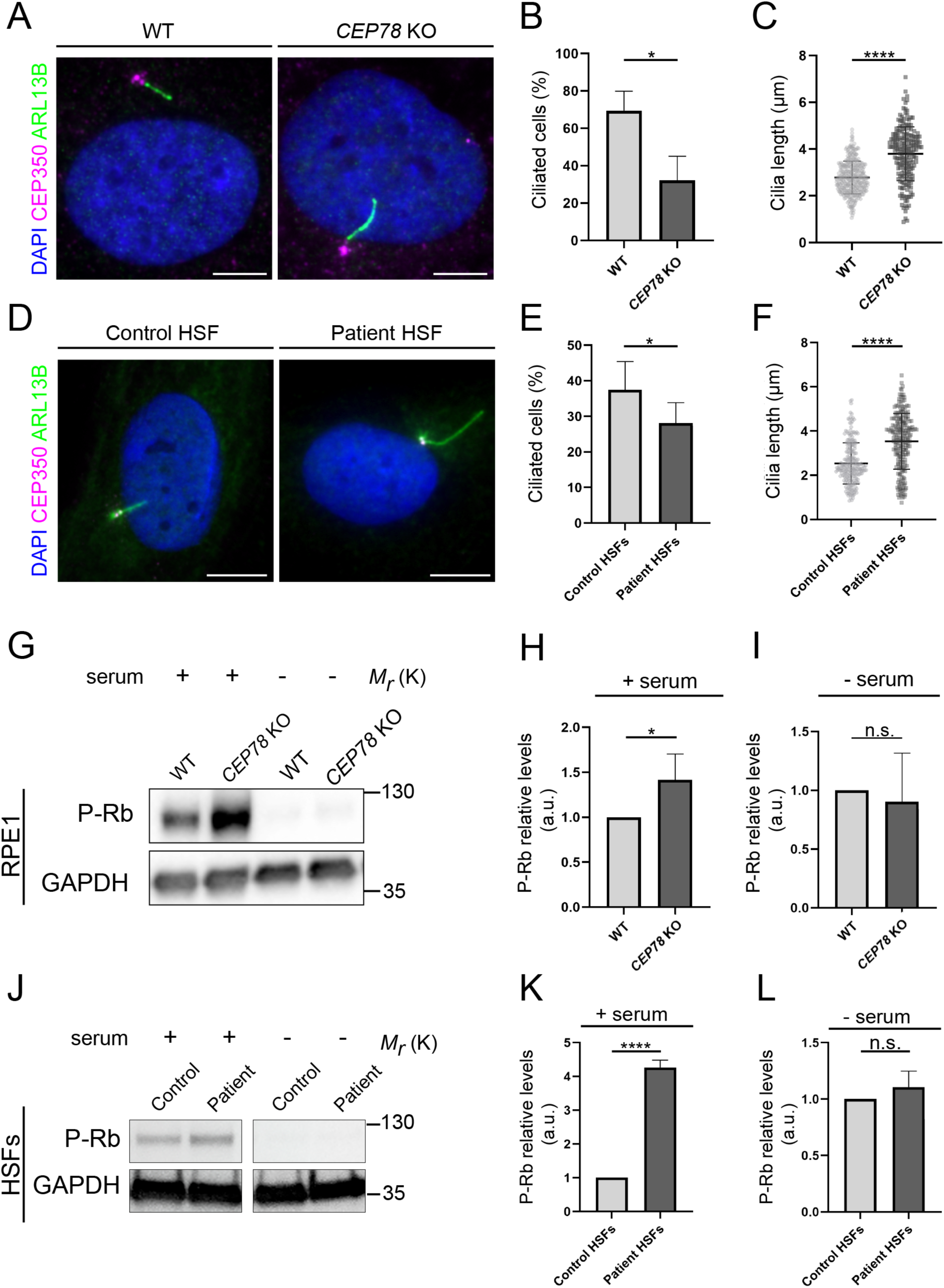
CEP78 mutant cells display fewer, but longer primary cilia. (**A**) Representative IFM images of serum-deprived RPE1 WT and *CEP78* KO cells stained with the indicated antibodies; DAPI marks the nucleus. Scale bar, 5μm. (**B**, **C**) Quantification of the percentage of ciliated cells (**B)** and the length of residual cilia (**C)** in WT and *CEP78* KO cells, based on images as shown in (A). Data were normalized in relation to WT values. Student’s t-test (two-tailed, unpaired) was used for the statistical analysis. Graphs in (B) represent accumulated data from three individual experiments (n=384 for WT and n=322 for *CEP78* KO cells). Graphs in (C) show data from three individual experiments (n=338 for WT and n=198 for *CEP78* KO cells). (**D**) Representative IFM images of serum-deprived control and *CEP78* mutant (Patient) HSFs stained with the indicated antibodies; DAPI marks the nucleus. Scale bar, 5 μm. (**E**, **F**) Quantification of the percentage of ciliated cells (**E**) and the length of residual cilia (**F**) in control and *CEP78* patient HSFs (data from HSFs derived from patient 2702r34, individual II-3 [29]), based on images as shown in (D). Graphs in (E) represent accumulated data from seven individual experiments (n= 678 for control HSFs; n= 707 for patient-derived HSFs). Graphs in (F) show data from seven individual experiments (n= 237 for control HSFs; n= 216 for patient-derived HSFs). Data were normalized in relation to control values. A Student’s t-test (unpaired, two-tailed) was performed to address differences in the ciliary frequency and length between control and patient HSFs. (**G**) Western blot analysis of Rb phosphorylated on S807/811 (P-Rb) in RPE1 WT and *CEP78* KO cells grown with or without serum. (**H**, **I**) Quantification of data shown in (G) from three independent experiments analyzed in duplicates. (**J**) P-Rb blots from HSFs derived from control and patient HSFs grown with or without serum (data from HSFs derived from patient F3: II:1; [30]). GAPDH was used as a loading control. (**K, L**) Quantification of data shown in (J) from three independent experiments analyzed in duplicates. Student’s t-test (two-tailed, unpaired) was used for the statistical analysis in (H, I) and (K, L). Error bars of graphs represent SD and data are shown as mean ± SD. a.u., arbitrary units; (*) p<0.05; (****) p<0.0001; n.s., not statistically significant.

In agreement with our observations in RPE1 cells, similar results were obtained using previously described patient-derived *CEP78*-deficient or control HSFs (Figure 1D-F) [29, 30]. The low ciliation frequencies of serum-deprived *CEP78* mutant cells were not secondary to cell cycle defects, as judged by western blot analysis with antibody against retinoblastoma protein phosphorylated at S807/811 (P-Rb; Figure 1G-L). However, in serum-fed cells, P-Rb levels were significantly higher in *CEP78* mutant cells compared to controls (Figure 1G-L), indicating that mutant cells may progress slower through S-phase [36]. This is in line with previous reports indicating that CEP78 protein levels are upregulated in late S-G2 phase suggesting a role for CEP78 at this cell cycle stage, e.g. during centriole duplication [27, 35]. We conclude that CEP78 deficient RPE1 and HSF cells display reduced ciliation frequencies as well as increased length of remaining cilia, thus reconciling previous observations [29, 30, 34].

### Analysis of RPE1 WT and *CEP78* KO cells expressing FLAG- or mNG-tagged CEP78 or CEP78^L150S^ fusions

To confirm the above results, we first set out to perform a rescue experiment by expressing FLAG-CEP78 in WT and *CEP78* KO RPE1 cells followed by serum-deprivation and IFM analysis using FLAG and ARL13B antibodies. In parallel, we performed similar experiments with FLAG-CEP78^L150S^. We first analyzed the localization and expression levels of the two FLAG fusions in transiently transfected serum-deprived RPE1 cells. In 4% paraformaldehyde-fixed cells we observed that the ARL13B antibody labeled the cilium itself as well as the basal body, but not the daughter centriole. The basal body pool of ARL13B likely corresponds to that present in the mother centriole-associated vesicle of mitotic centrosomes [37]. The transiently expressed FLAG-CEP78 fusion protein localized to both centrioles at the base of cilia as expected [27, 29, 35], whereas centriolar localization of FLAG-CEP78^L150S^ was severely compromised although not completely abolished (Figure 2A, B). The latter result is in line with previous reports showing that the LRR region in the N-terminus of CEP78, which encompasses residue L150, is important for its localization to centrioles [27, 35]. Both FLAG fusions were expressed at similar levels in the cells (Figure 2-figure supplement 1); this was somewhat surprising given that endogenous CEP78^L150S^ is largely undetectable by western blot analysis of patient-derived HSFs suggesting its stability is compromised [30]. Therefore, we tested if the L150S mutation might affect binding of the CEP78 antibody used in [30] to CEP78. However, FLAG immunoprecipitation (IP) and western blot analysis of 293T cells expressing FLAG-CEP78 or FLAG-CEP78^L150S^ indicated that the CEP78 antibody binds equally well to WT CEP78 and the CEP78^L150S^ mutant protein. This analysis also revealed that endogenous CEP78 co-IPs with FLAG-CEP78 indicating that CEP78 can form homodimers/oligomers (Figure 2-figure supplement 2). These results indicate that the L150S mutation compromises the recruitment of CEP78 to the centrosome, but not its short-term stability, at least in RPE1 cells. We cannot rule out that the N-terminal FLAG tag may stabilize FLAG-CEP78^L150S^ and prevent it from being degraded.

**Figure 2.**
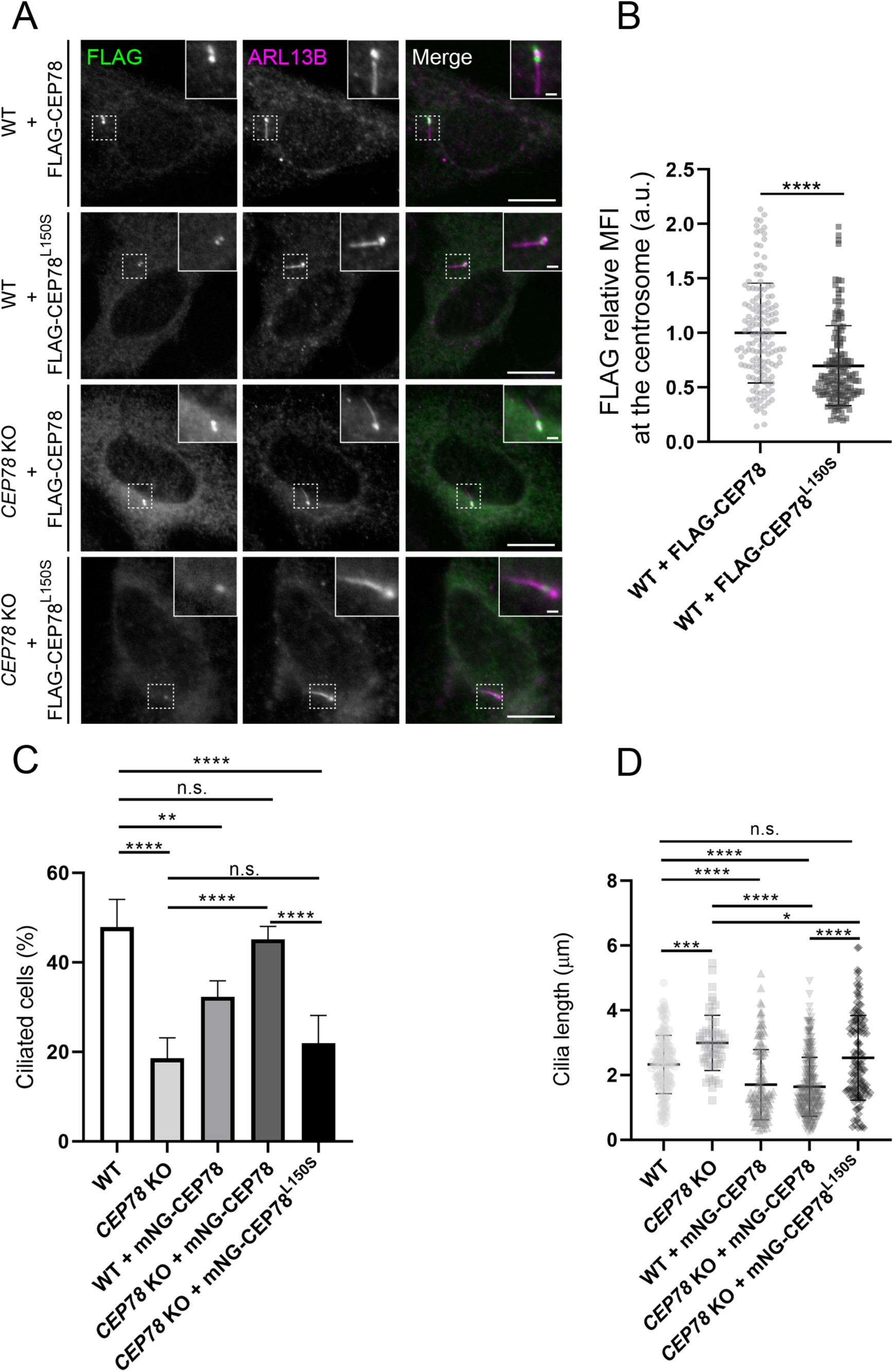
Analysis of RPE1 WT and *CEP78* KO cells expressing FLAG- or mNG-tagged CEP78 or CEP78^L150S^ fusions. (**A**) Representative IFM images of serum-starved WT and *CEP78* KO cells transiently expressing FLAG-CEP78 and FLAG-CEP78^L150S^ labeled with antibodies against FLAG (green) and ARL13B (magenta). Insets show enlarged views of the cilia-centrosome region. Scale bars: 5 μm in original images and 1 μm in magnified images. (**B**) Quantification of the relative MFI of FLAG-CEP78 and FLAG-CEP78^L150S^ at the centrosome in serum-starved WT cells. Accumulated data from three individual experiments (n=145 and n=151 for cells transfected with FLAG-CEP78 and FLAG-CEP78^L150S^, respectively). Student’s t-test (two-tailed, unpaired) was used for the statistical analysis. Data is shown as mean ±SD. (**C, D**). Quantification of the percentage of ciliated cells (**C**) or ciliary length (**D**) in serum-deprived WT and *CEP78* KO cells stably expressing mNG-CEP78 or mNG-CEP78^L150S^. Analysis on cilia numbers and length was performed through accumulated data from three independent experiments for WT, *CEP78* KO and WT + mNG-CEP78 cells and five independent experiments for *CEP78* KO cells expressing either the mNG-CEP78 or the mNG-CEP78^L150S^ (for ciliary frequency: n=304 for WT; n=322 for *CEP78* KO; n=425 for WT + mNG-CEP78; n=621 for C*EP78* KO + mNG-CEP78, and n=638 for C*EP78* KO + mNG-CEP78^L150S^; for ciliary length: n=145 for WT; n=56 for *CEP78* KO; n=138 for WT + mNG-CEP78; n=280 for C*EP78* KO + mNG-CEP78 and n=141 for C*EP78* KO + mNG-CEP78^L150S^). Ordinary one-way ANOVA with Dunnett’s multiple comparison test was used for the statistical analysis of the ciliary frequency and length amongst all groups, in relation to the mean of WT cells, designated as the control group. Differences between *CEP78* KO and *CEP78* KO+ mNG-CEP78; *CEP78* KO and *CEP78* KO + mNG-CEP78^L150S^ and *CEP78* KO + mNG-CEP78 and *CEP78* KO + mNG-CEP78^L150S^ were addressed in a pairwise fashion using an unpaired and two-tailed Student’s t-test. Error bars in (C) indicate SD and data in (D) is presented as mean ± SD. (*) p<0.05; (**) p<0.01; (***) p<0.001; (****) p<0.0001; n.s, not statistically significant; a.u., arbitrary units.

Next, we tested if FLAG-CEP78 or FLAG-CEP78^L150S^ could rescue the ciliary phenotypes of *CEP78* KO cells. However, upon transient expression of the FLAG-CEP78 fusions we could not rescue the cilia frequency and length phenotype of the *CEP78* KO cells, and cilia frequency and length were also affected in WT control cells transiently expressing these fusions (data not shown). We therefore generated cell lines stably expressing mNeonGreen (mNG)-tagged CEP78 or CEP78^L150S^ at ca. 3-4 times the endogenous CEP78 level by viral transduction of mNG-CEP78 constructs into WT and *CEP78* KO RPE1 cells, respectively (Figure 2-figure supplement 3A). Microscopic examination of these lines indicated that mNG-CEP78 was concentrated at the centrosome as expected, whereas mNG-CEP78^L150S^ was diffusely located in the cytosol (Figure 2-figure supplement 3B), in agreement with results for transiently expressed FLAG-CEP78/CEP78^L150S^ fusions (Figure 2A, B). Importantly, stably expressed mNG-CEP78 could fully rescue the ciliation frequency phenotype of *CEP78* KO cells whereas mNG-CEP78^L150S^ could not (Figure 2C). In addition, we observed that WT cells stably expressing mNG-CEP78 have significantly reduced ciliation frequency compared to untransfected WT cells (Figure 2C), indicating that mild overexpression of mNG-CEP78 seems to inhibit ciliogenesis. Furthermore, WT or *CEP78* KO cells stably expressing mNG-CEP78 had significantly shorter cilia than untransfected WT and *CEP78* KO cells, or *CEP78* KO cells expressing mNG-CEP78^L150S^. However, we note that cilia length in the latter was significantly shorter than untransfected *CEP78* KO cells (Figure 2D). Together these results suggest that mNG-CEP78 is able to fully rescue the cilia frequency phenotype of *CEP78* KO cells whereas mNG-CEP78^L150S^ is not. Furthermore, both mNG-CEP78 and mNG-CEP78 ^L150S^ promote ciliary shortening, but mNG-CEP78 does it more efficiently than mNG-CEP78 ^L150S^. We conclude that the observed ciliary frequency and length phenotypes of the *CEP78* KO cells are caused specifically by CEP78 loss, and that the L150S mutations impairs CEP78 centrosome localization and function.

### CEP78 interacts with the EDD1-DYRK2-DDB1^VPRBP^ complex and CEP350

To explore how CEP78 is recruited to centrioles to regulate ciliary frequency and length we first used Stable Isotope Labeling by Amino acids in Cell culture (SILAC)-based quantitative mass spectrometry (MS) to identify the differential interactomes of FLAG-CEP78 and FLAG-CEP78^L150S^ expressed in 293T cells (see Materials and methods for details). We identified interaction between CEP78 and several components of the EDD1-DYRK2-DDB1^VPRBP^ complex (Figure 3A), in agreement with previous work showing direct binding of CEP78 to VPRBP within this complex [27]. Interestingly, the MS results indicated that the CEP78^L150S^ mutation dramatically reduces this interaction (Figure 3A), which we confirmed by IP and western blot analysis of 293T cells co-expressing Myc-VPRBP and FLAG-CEP78 or FLAG-CEP78^L150S^ (Figure 3B). In addition, our interactome analysis identified CEP350 as potential novel CEP78 binding partner (Figure 3A). Using IP and western blot analysis in 293T cells co-expressing FLAG-CEP78 or FLAG-CEP78^L150S^ with a Myc-tagged N-terminal or C-terminal CEP350 truncation we found that CEP78 binds to the N-terminal region of CEP350 (residues 1-983; Figure 3C, E); reciprocal co-IP with Myc-tagged CEP350 constructs confirmed this result (Figure 3-figure supplement 1A). Furthermore, we found that the interaction between CEP78 and CEP350 N-terminus is reduced by the CEP78^L150S^ mutation (Figure 3C). Using a similar approach, we could not detect interaction between CEP78 and endogenous FOP (Figure 3C), whereas under similar conditions endogenous FOP was readily co-IPed with the C-terminus of CEP350 (Figure 3-figure supplement 1A), in agreement with a previous report [38]. We also mapped the CEP350 interaction site in CEP78 and found that CEP350 primarily binds to the C-terminal region of CEP78 comprising residues 395-722 (Figure 3D, E), although a weak interaction with a fragment comprising the entire LRR region (residues 2-403) was also observed. In contrast, truncation of the latter fragment into two smaller fragments abolished binding to the CEP350 N-terminus (Figure 3D, E). Binding of CEP78 to VPRBP similarly requires the entire LRR region of CEP78 [27] and we therefore wondered whether interaction between CEP350 and CEP78 might be mediated by VPRBP. However, Myc IP experiments and western blot analysis failed to reveal physical interaction between VPRBP and Myc-CEP350 N-terminus (Figure 3, figure supplement 1A). Moreover, GFP IP of cells co-expressing GFP-CEP78 and Myc-CEP350 N- or C-terminal fragments showed that GFP-CEP78 can bind to Myc-CEP350 N-terminus and endogenous VPRBP at the same time, indicating that CEP350 does not compete with VPRBP for binding to CEP78 (Figure 3, figure supplement 1B). We conclude that CEP78 binds independently to VPRBP and the N-terminus of CEP350 and that both of these interactions are reduced by the CEP78^L150S^ mutation.

**Figure 3.**
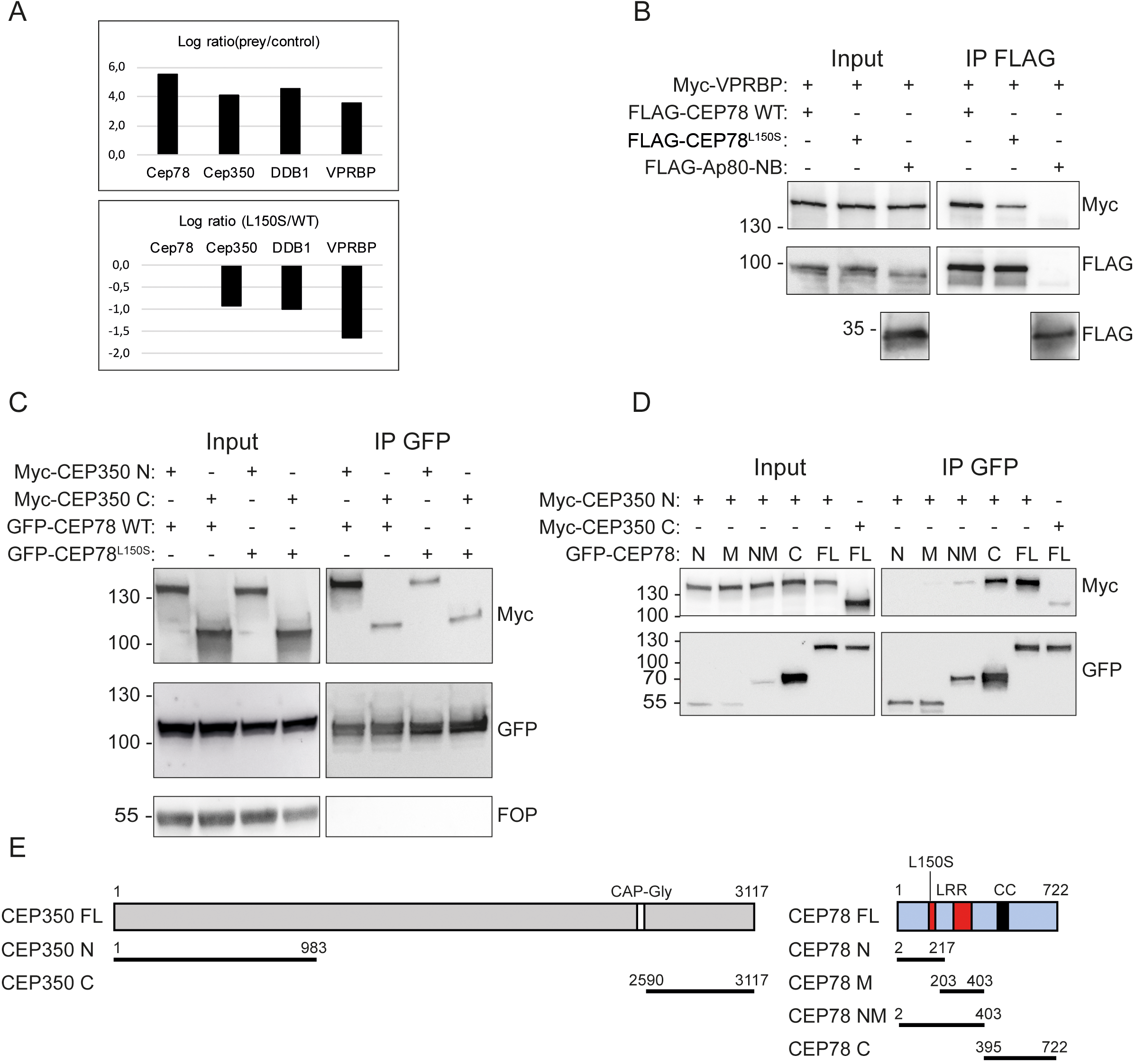
CEP78 interacts with the EDD-DYRK2-DDB1^VPRBP^ complex and CEP350. (**A**) Overview of results from CEP78 interactome analysis in 293T cells grown in SILAC medium. Cells expressing FLAG-CEP78 WT, FLAG-CEP78^L150S^ or FLAG-Ap80 (negative control) were subjected to FLAG IP and pellets analyzed by mass spectrometry. The upper panel displays affinity enrichment of Cep78 and interaction partners in FLAG-CEP78 WT cells relative to control cells. The lower panel shows how the CEP78^L150S^ mutation weakens these interactions relative to CEP78 WT using ratios normalized for equal affinity enrichment of FLAG-tagged CEP78. (**B**-**D**) 293T cells expressing the indicated Myc-, FLAG-, or GFP-fusions were subjected to IP with anti-FLAG (**B**) or -GFP (**C**, **D**) beads and input and pellet fractions analyzed by SDS-PAGE and western blotting with Myc, FLAG or GFP antibodies as indicated. FLAG-Ap80-NB used in (B) is a negative control. Molecular mass markers are indicated in kDa to the left of the blots. (**E**) Schematic of the CEP350 and CEP78 fusions used in IP analysis. LRR, leucine-rich repeat; CC, coiled coil. The CEP350 constructs were described in [54, 58]; the CEP78 constructs were generated in this study based on published sequence information [27, 30, 35]. N, N-terminal region; M, middle region; NM, N-terminal plus middle region; C, C-terminal region; FL, full length.

### CEP350 regulates the stability and centrosomal recruitment of CEP78

In non-ciliated HeLa and U2OS cells endogenous CEP78 was reported to localize to both centrioles during all phases of the cell cycle and appeared to concentrate at the centriole wall [35]. In ciliated RPE1 cells CEP78 was shown to localize to the distal end of both centrioles, with a preference for the mother centriole/basal body [27]. Similarly, CEP350 was reported to localize to the distal end of the mother centriole/basal body, near the subdistal appendages, in ciliated RPE1 cells [24, 26, 39]. In agreement with these reports and with our interactome and IP results (Figure 3), 3D Structured Illumination Microscopy (SIM) showed partial co-localization of mNG-CEP78 with CEP350 at the distal end of the centrioles in RPE1 cells, adjacent to the distal appendage marker CEP164 and the centriole capping complex protein CP110 (Figure 4A). Superimposing fluorescence images on electron micrographs of the mother centriole [40] indicates that mNG-CEP78 localizes to a ring-structure of centriole-sized diameter at the distal half of the centriole, encompassed by a wider unclosed ring-structure of CEP350. mNG-CEP78 but not CEP350 depicted decreased levels at the daughter centriole as compared to the mother. Furthermore, using a previously characterized RPE1 *CEP350* KO cell line [24] that displays significantly reduced centrosomal levels of CEP350 (Figure 4-figure supplement 1), we found that depletion of CEP350 not only reduces localization of CEP78 to the centrosome, but also its total cellular level (Figure 4B-G). Similarly, we found significantly reduced levels of CEP78 in *FOP* KO cells (Figure 4 B, C), in line with the decreased centrosomal levels of CEP350 observed in these cells [24]. These results indicate that CEP350 promotes the recruitment of CEP78 to the mother centriole/basal body as well as its overall stability.

**Figure 4.**
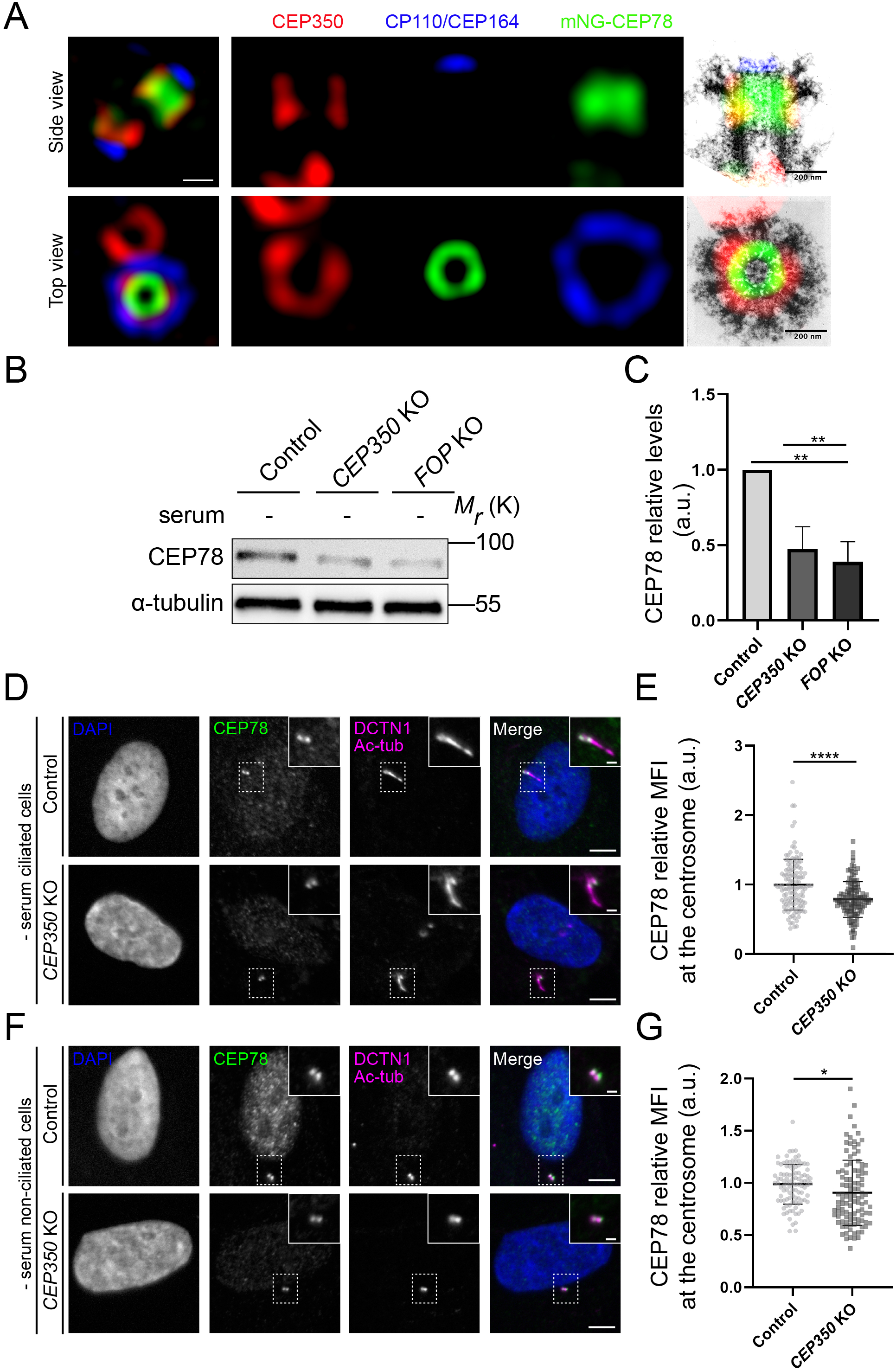
CEP350 regulates stability and centrosomal recruitment of CEP78. (**A**) 3D-SIM analysis of mNG-CEP78 and CEP350 localization in WT RPE1 cells, relative to CP110 and CEP164 (maximal z-projections). Cells were fixed and stained with the indicated antibodies. Montage panels show single channel images of the mother centriole region (left panels) and overlays on previously published electron micrographs of isolated mother centrioles from KE37 cells, reproduced with permission from [40] (top: cross-sections of proximal and subdistal-appendage region; bottom: side-view). Scale bar, 200 nm. (**B**) Western blot analysis of CEP78 and *α*-tubulin (loading control) in serum-deprived hTERT-RPE1-BFP-Cas9 control (pMCB306), *CEP350* KO or *FOP* KO cells [24]. (**C**) Quantification of data in (B), based on three independent experiments analyzed in duplicates. Error bars indicate SD. Statistical analysis was done using Student’s t-test (unpaired, two-tailed). (**D, F**) IFM analysis of serum-deprived control and *CEP350* KO ciliated (D) and non-ciliated cells (F) using antibodies as indicated. Dashed boxes indicate cropped images to highlight the centrosomal/ciliary area. Scale bars: 5 µm in representative images and 1 µm in insets. (**E, G**) Quantification of the relative MFI of CEP78 at the centrosome in serum-starved, ciliated (E) and non-ciliated (G) control and *CEP350* KO cells, based on images as shown in (D) and (F), respectively. Statistical analysis was done using a two-tailed and unpaired Mann-Whitney test for ciliated cells and two-tailed and unpaired Student’s t-test for non-ciliated cells. Accumulated data from three individual experiments (n=128 and 132 for control and *CEP350* KO ciliated cells, respectively; n= 101 and 107 for control and *CEP350* KO non-ciliated cells, respectively). a.u., arbitrary units; (*) p<0.05; (**) p<0.01; (****) p<0.0001.

### CEP78 mutant cells display reduced centrosomal levels of EDD1

Next, we asked if CEP78 might affect the centrosomal recruitment of components of the EDD1-DYRK2-DDB1^VPRBP^ complex and/or CEP350, and analyzed the cellular or centrosomal levels of relevant complex components in RPE1 WT and *CEP78* KO cells. The results showed that serum-deprived, ciliated *CEP78* KO cells display significantly increased centrosomal levels of CEP350 (Figure 5A, B) as compared to controls. However, in serum-deprived, non-ciliated cells, the centrosomal levels of CEP350 were not significantly different in *CEP78* KO cells compared to WT controls (Figure 5C, D). Similar results were obtained for CEP350 in control and CEP78 deficient patient HSFs (Figure 5E-H). We also analyzed the cellular levels of VPRBP by western blotting and the centrosomal levels of VPRBP and EDD1 by IFM in RPE1 WT and *CEP78* KO cells. In agreement with a previous study [27] we found that cellular and centrosomal levels of VPRBP were not significantly reduced in serum-fed or serum-deprived *CEP78* KO cells compared to controls (Figure 6A-C; Figure 6-figure supplement 1). Thus, CEP78 seems to be dispensable for recruitment of VPRBP and CEP350 to the centrosome. In contrast, centrosomal levels of EDD1 were significantly reduced in RPE1 *CEP78* KO cells compared to WT controls, both in ciliated and non-ciliated cells (Figure 6D-G). Thus, CEP78 is required for efficient recruitment of EDD1 to the centrosome. To corroborate this result, we measured the centrosomal levels of EDD1 in RPE1 *CEP350* KO cells and found that EDD1 levels were also significantly reduced when compared to the controls (Figure 6-figure supplement 2). Together our data indicate that CEP350, by recruiting CEP78 to the centrosome, also promotes the centrosomal recruitment of EDD1.

**Figure 5.**
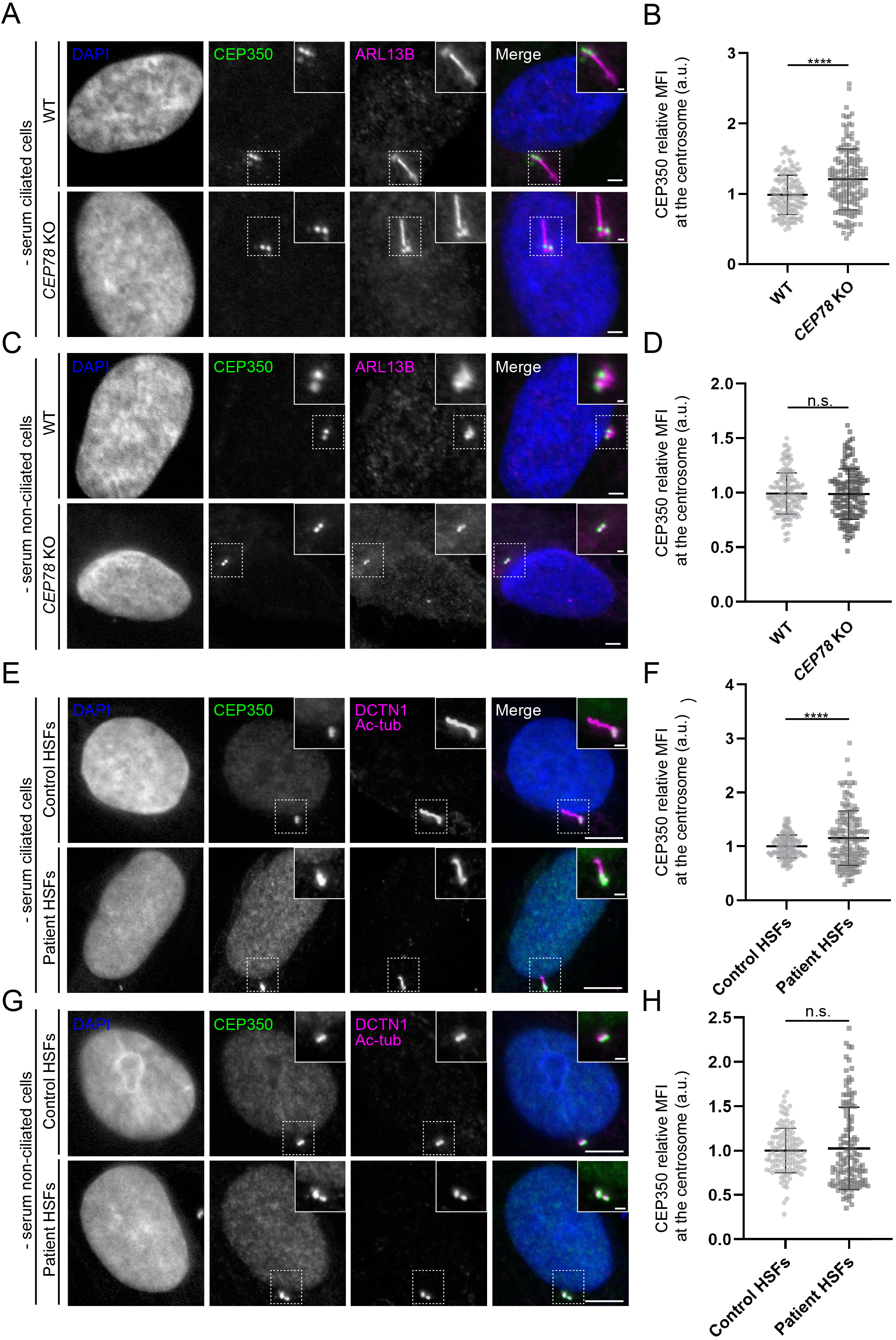
Altered centrosomal levels of CEP350 in *CEP78* mutant cells. (**A, C, E, G**) Representative IFM images of ciliated (**A**) and non-ciliated (**C**) serum-deprived RPE1 WT and *CEP78* KO cells, and ciliated (**E**) and non-ciliated (**G**) serum-deprived control and CEP78 deficient (Patient) HSFs (combined data from HSFs from patient F3: II:1 [30] and patient 2702r34, individual II-3 [29]) labeled with antibodies indicated in the figure. Insets show closeups of the centrosomal/ciliary area (dashed boxes). Scale bars: 5 µm in representative images and 1 µm in closeups. (**B, D**). Quantification of CEP350 relative MFI at the centrosome based on images as shown in (A, C). Data is shown as mean ± SD. A Student’s t-test was used as statistical analysis between the two groups based on three individual experiments (n=157 or 158 for ciliated cells and n=144-152 for non-ciliated cells). (**F, H)** Quantification of CEP350 relative MFI at the centrosome in HSFs based on images as shown in (E, G). Data is shown as mean ± SD. Student’s t-test was used as statistical analysis between the two cell groups based on seven individual experiments with a total of 154 and 156 analyzed ciliated cells (**F**), and three individual experiments with a total of 122 or 126 analyzed non-ciliated cells (**H**). a. u., arbitrary units; n.s, not statistically significant; (****) p<0.0001.

**Figure 6.**
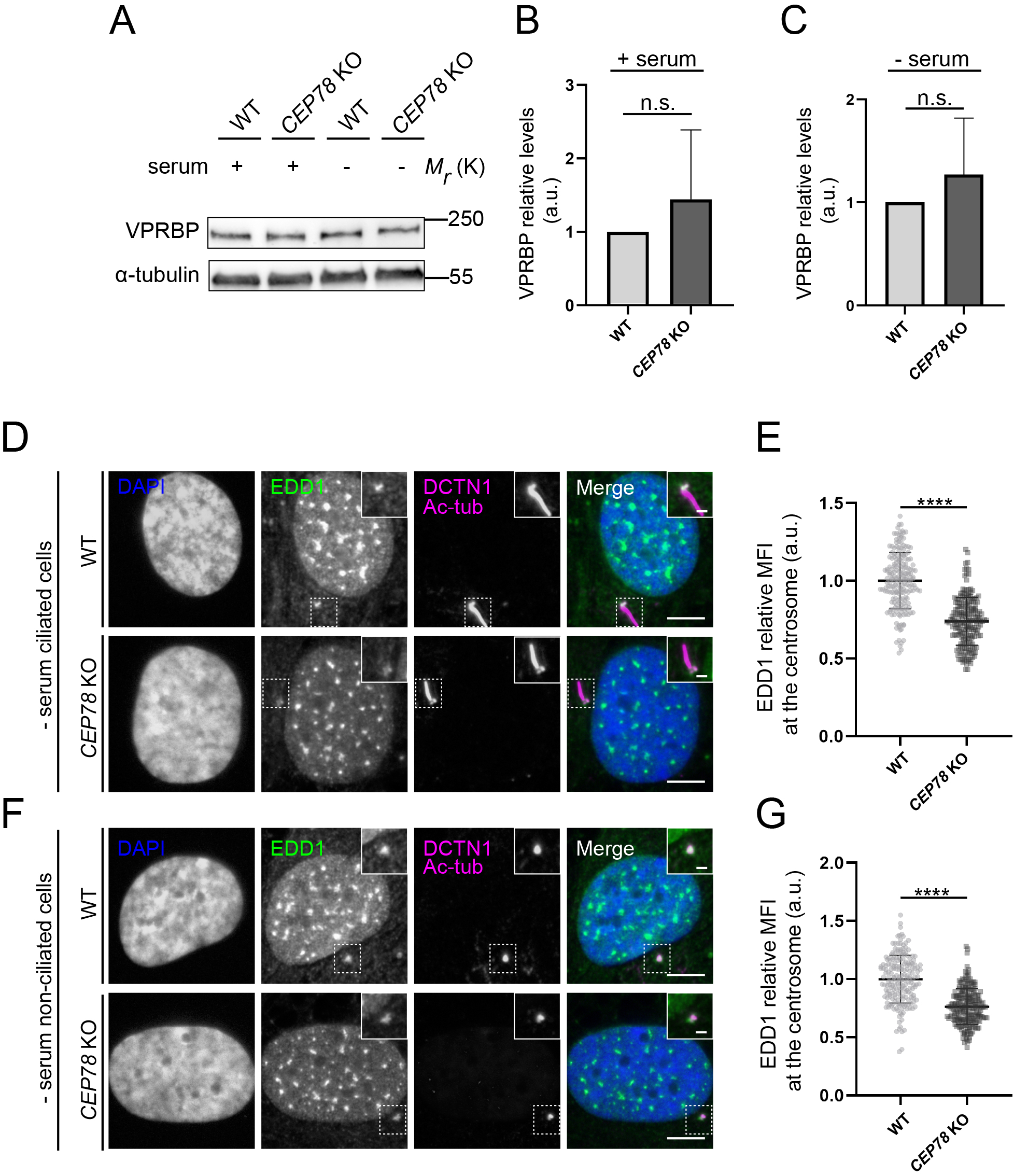
Analysis of EDD-DYRK2-DDB1^VPRBP^ complex components in CEP78 deficient cells. (**A**) Western blot analysis of VPRBP in lysates from serum-fed and serum-deprived RPE1 WT and *CEP78* KO cells. α-tubulin was used as a loading control (**B, C**) Quantification of the VPRBP relative levels in the different conditions depicted in (A). Statistical analysis was performed using a Student’s t-test (unpaired, two-tailed) from five independent experiments analyzed in duplicates. Error bars indicate SD. (**D, F)** Representative IFM images of ciliated (**D**) and non-ciliated (**F**) serum-deprived RPE1 WT and *CEP78* KO cells labeled with antibodies against EDD1 (green) and DCTN1 plus acetylated tubulin (magenta). DAPI was used to mark the nucleus (blue). Insets show enlarged views of the cilium-centrosome region. Scale bars: 5 μm in original images and 1 μm in closeups. (**E, G)** Quantification of the EDD1 MFI at the centrosome based on images as shown in (D) and (F) using a two-tailed and unpaired Student’s t-test. Data is shown as mean ± SD. Student’s t-test from three independent experiments (n=194 and n=201 for ciliated WT and *CEP78* KO cells, respectively; n=194 and n=217 for non-ciliated WT and *CEP78* KO cells, respectively). Data is shown as mean ± SD. a.u., arbitrary units; n.s., not statistically significant; (****) p<0.0001.

### Increased cellular and centrosomal levels of CP110 in CEP78 deficient cells

Since EDD1 is an E3 ubiquitin-protein ligase that negatively regulates CP110 levels [27], and whose depletion impairs ciliogenesis [41, 42], we asked if the loss of CEP78 would lead to altered levels of CP110 at the centrosome. Indeed, western blot and IFM analysis of RPE1 WT and *CEP78* KO cells revealed that total cellular and centrosomal CP110 levels are significantly increased in *CEP78* KO cells compared to WT controls, both under serum-deprived (Figure 7A-D, F, G; Figure 7-figure supplement 1A, B) and serum-fed conditions (Figure 7-figure supplement 2A-D), whereas CP110 levels appeared normal upon stable expression of mNG-CEP78 in the *CEP78* KO background (Figure 7-figure supplement 1C). We also analyzed CP110 levels specifically at the mother centriole of ciliated RPE1 WT and *CEP78* KO cells and found no significant differences between them (Figure 7E). Similar results were obtained using *CEP78* mutant and control HSF cells (Figure 7G-H). Since a previous study reported that siRNA-mediated depletion of CEP78 in RPE1 and HeLa cells reduces the cellular levels of CP110 [27], a result contradicting our own observations (Figure 7; Figure 7-figure supplement 1 and 2), we tested if the elevated CP110 levels seen in our *CEP78* mutant cells might be due to compensatory upregulation of CP110 mRNA expression. However, RNA-Seq analysis indicated no significant changes in CP110 mRNA levels in patient-derived, *CEP78* deficient HSF cultures compared to controls; mRNA levels of CEP350, VPRBP and DDB1 in *CEP78* deficient HSF cultures were also similar to those of controls (Figure 7-figure supplement 3). Furthermore, a cycloheximide chase experiment indicated that CP110 is more stable upon serum deprivation of *CEP78* KO cells compared to WT controls (Figure 7-figure supplement 4). Taken together, we conclude that loss of CEP78 leads to elevated cellular and centrosomal levels of CP110 and that this is likely due to increased stability of CP110. Furthermore, our results indicate that in ciliated *CEP78* KO cells, the amount of CP110 present specifically at the mother centriole is not significantly higher than in control cells.

**Figure 7.**
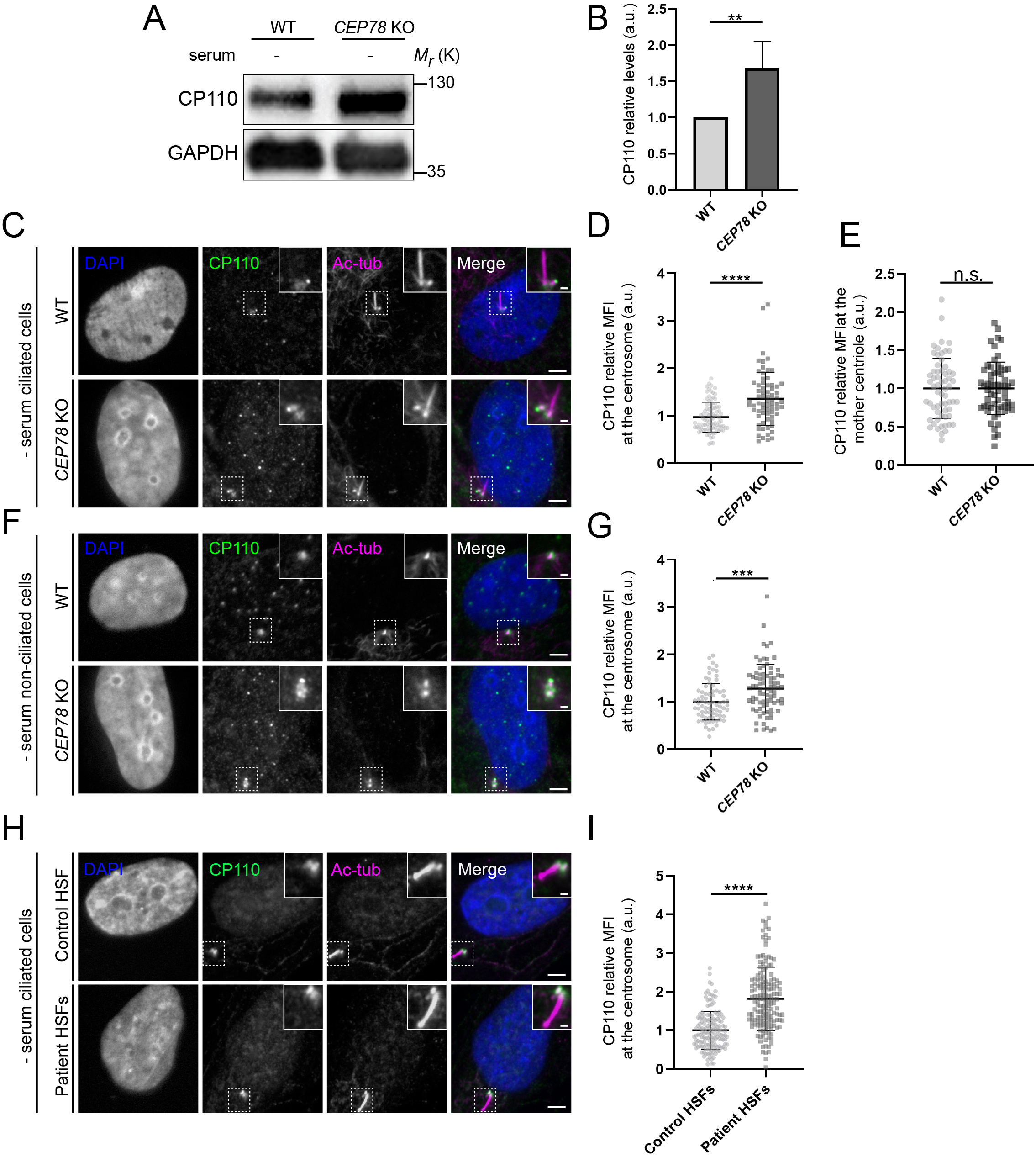
CEP78 deficient cells display elevated cellular and centrosomal levels of CP110. (**A**) Western blot analysis of lysates from serum-deprived RPE1 WT and *CEP78* KO cells using indicated antibodies. GAPDH was used as a loading control. (**B**) Quantification of the data shown in (A), based on four independent experiments analyzed in duplicates. Error bars indicate SD. Statistical analysis was done using Student’s t-test (unpaired, two-tailed). (**C, F**) Representative IFM images of serum-starved ciliated (**C**) and non-ciliated (**F**) RPE1 WT and *CEP78* KO cells labeled with antibodies against CP110 (green), combined DCTN1 and acetylated tubulin (Ac-tub; magenta) and DAPI to mark the nucleus (blue). Insets show enlarged views of the cilium-centrosome area. Scale bars: 5 µm in original images and 1 µm in closeups. (**D, G**) Quantification of the relative MFI of CP110 at the centrosome based on images as shown in (C) and (F), respectively. Student’s t-test (unpaired, two-tailed) from three independent biological experiments (n= 90 and n=75 for ciliated RPE1 WT and *CEP78* KO cells, respectively; n=75 and n=82 for non-ciliated RPE1 WT and CEP78 KO cells, respectively) was used for statistical analysis. Data is presented as mean ± SD. (**E**) Quantification of the relative MFI of CP110 at the mother centriole based on images shown in (C) Mann Whitney test (two-tailed and unpaired) was used as statistical analysis based on two independent experiments (n=61 and n=62 for RPE1 WT and CEP78 KO cells, respectively). Data is presented as mean ± SD. (**H**) Representative IFM images of serum-deprived healthy and CEP78 deficient (Patient) HSFs (data from patient 2702r34, individual II-3 described in [29]) labeled with the indicated antibodies. DAPI was used as counterstaining to mark the nucleus. Dashed lines show closeup images of the centrosome region. Scale bars: 5 µm in original images and 1 µm in closeups. (**I**) Quantification based on observations of 151 and 155 cells of CEP78 control and patient cells, respectively, from three individual experiments. Student’s t-test (unpaired and two-tailed) was used to assess differences between the two groups. a.u., arbitrary units; n.s., not statistically significant; (**) p<0.01; (***) p<0.001; (****) p<0.0001.

### Partial depletion of CP110 normalizes ciliation frequency of *CEP78* KO cells

CP110 is a key negative regulator of ciliogenesis [5, 43] that was also shown to be required for ciliogenesis initiation as well as ciliary length control [44, 45]. The above results therefore prompted us to test if the reduced ciliation frequency and/or increased length of remaining cilia observed in *CEP78* deficient cells could result from elevated centrosomal CP110 levels in these cells. When we used siRNA to partially deplete cellular and centrosomal CP110 from RPE1 WT and *CEP78* KO cells and subjected the cells to serum deprivation (Figure 8A, B; Figure 8-figure supplement 1), we found that the *CEP78* KO cells restored their ciliation frequency to WT levels, whereas partial CP110 depletion had little effect on cilia numbers in WT cells grown under similar conditions (Figure 8C, D). In contrast, depletion of CP110 from *CEP78* KO cells did not rescue the ciliary length defect of residual cilia; these cilia were still significantly longer than those of WT controls (Figure 8C, E). Thus, partial depletion of CP110 from *CEP78* KO cells can rescue their reduced ciliation frequency phenotype but not the increased length of remaining cilia. This result is in line with our observation that a *CEP78* KO clone expressing a truncated version of CEP78 (clone #44) displays significantly reduced ciliation frequency, but normal length of the cilia that do form (Figure 1-figure supplement 1), implying that CEP78 regulates ciliogenesis and ciliary length by separate mechanisms.

**Figure 8.**
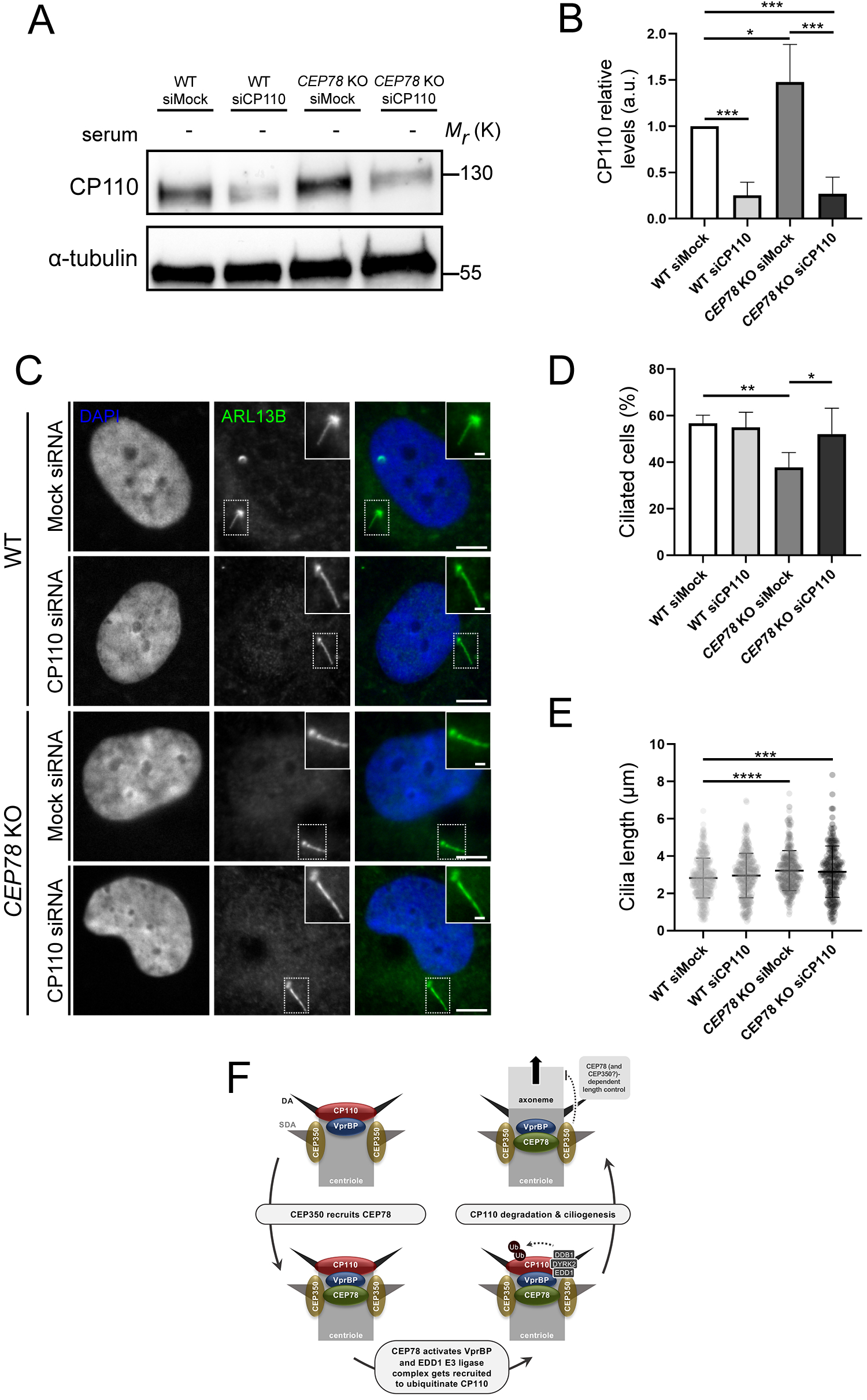
Depletion of CP110 from *CEP78* KO cells rescues their ciliation frequency phenotype. (**A**) Representative western blots of RPE1 WT or *CEP78* KO cells treated with control (siMock) or CP110-specific siRNA (siCP110). Cells were deprived of serum and blots were probed with antibody against CP110 or *α*-tubulin (loading control). (**B**) Quantification of relative cellular CP110 levels under different conditions based on blots as shown in (A) (n=5 for WT and n=4 for *CEP78* KO cells; each sample was analyzed in duplicates). Error bars indicate SD. Data was normalized in relation to the WT cells transfected with Mock siRNA (WT siMock), designated as the control group. Differences of CP110 relative levels between *CEP78* KO cells transfected with Mock or CP110 siRNA and WT cells transfected with CP110 siRNA in relation to control were assessed by performing an ordinary one-way ANOVA with a Dunnet’s multiple comparison test. Differences of CP110 cellular levels between *CEP78* KO cells transfected with the same siRNAs was addressed using an unpaired and two-tailed Student’s t-test. (**C**) Representative IFM images of serum-deprived RPE1 WT and *CEP78* KO cells treated with control (siMock) or CP110-specific siRNA (siCP110) and stained for ARL13B (green). DAPI marks the nucleus. Dashed boxes indicate cropped images to highlight the centrosomal/ciliary zone. Scale bars, 5 μm in representative images and 1µm in insets. (**D**, **E**) Quantification of percentage of ciliated cells (**D**) and length of remaining cilia (**E**) in RPE1 WT or *CEP78* KO cells treated with control (siMock) or CP110-specific siRNA (siCP110). The quantification is based on images as shown in (C) and represents data from four and five independent experiments, respectively, for RPE1 WT (n=554-617cells per condition) and *CEP78* KO cells (n=846-856 cells per condition). Ordinary one-way ANOVA with Dunnett’s multiple comparison test was used, to compare the mean of the remaining groups in relation with the mean of RPE1 WT cells treated with siMock, which was designated as the control group. Differences between *CEP78* KO cells transfected with the above-mentioned siRNA were discriminated using an unpaired and two-tailed Student’s t-test. Bars in (D) represent SD and data in (E) is presented as mean ± SD. a.u., arbitrary units; (*) p<0.05; (**) p<0.01; (***) p<0.001; (****) p<0.0001. (**F**) Proposed model for how CEP78 regulates ciliogenesis and ciliary length control. See text for details.

## Discussion

Several studies have shown that loss-of-function mutations in *CEP78* are causative of ciliopathy characterized by CRDHL [29–32], and that cells lacking CEP78 display reduced ciliation frequency [34] or abnormally long cilia [29, 30]. However, the molecular mechanisms by which CEP78 regulates cilium biogenesis and length were so far unclear. In this study, we found that loss of CEP78 leads to decreased ciliation frequency as well as increased length of the residual cilia that do form, both in RPE1 cells and patient-derived HSFs. Recue experiments in *CEP78* KO RPE1 cells stably expressing mNG-CEP78 or mNG-CEP78^L150^ supported that the observed phenotypes were specifically caused by loss of CEP78 at the centrosomes. Interestingly, we found that both ciliated and non-ciliated CEP78 deficient cells fail to efficiently recruit EDD1 to the centrosome, whereas cellular and overall centrosomal levels of CP110 are dramatically increased. However, in the subpopulation of *CEP78* KO cells that are ciliated, CP110 levels specifically at the mother centriole were similar to those of ciliated control cells, implying that the elevated centrosomal CP110 level observed in these cells is caused by a pool of CP110 located at the daughter centriole and/or pericentriolar area. This result also indicates that the abnormally long cilia phenotype seen in the *CEP78* KO cells is not caused by increased CP110 levels at the mother centriole. Supportively, siRNA-mediated depletion of CP110 could restore cilia frequency, but not length, to normal in the *CEP78* KO cells. Furthermore, a *CEP78* mutant line, clone #44, expressing a shorter version of CEP78 likely lacking the N-terminus displayed reduced ciliation frequency but normal length of residual cilia, substantiating that CEP78 regulates ciliogenesis and cilia length by distinct mechanisms. Specifically, our results indicate that CEP78 promotes ciliogenesis by negatively regulating CP110 levels at the mother centriole via activation of VprBP, but negatively regulates ciliary lengthening independently of CP110 (Figure 8F).

The removal of CP110 from the distal end of the mother centriole constitutes a key step in the initiation of ciliogenesis and is regulated by a growing number of proteins that include TTBK2 [19] and components of the UPS system such as Neurl-4 [46] and the cullin-3 (CUL3)–RBX1–KCTD10 complex [47]. The EDD1-DYRK2-DDB1^VPRBP^ complex was similarly shown to mediate degradation of CP110 and to interact with CEP78 [27], but the precise consequences of these activities for ciliogenesis were unclear. Our work strongly suggests that CEP78 functions together with the EDD1-DYRK2-DDB1^VPRBP^ complex to negatively regulate centrosomal CP110 levels at the onset of ciliogenesis, leading to its removal from the distal end of the mother centriole (Figure 8F). This is in contrast to a previously proposed model, which suggested that CEP78 inhibits degradation of CP110 by the EDD1-DYRK2-DDB1^VPRBP^ complex [27]. The reason for this discrepancy is unclear, but may be due to different experimental approaches used in our study and the paper by Hossain et al. [27]. In this latter study, siRNA was used to partially deplete CEP78 from cultured cells, whereas we performed our experiments on cells with endogenous *CEP78* mutations. CP110 removal from the distal end of the mother centriole occurs concomitantly with docking of the centriole to vesicles or the plasma membrane at the onset of ciliogenesis [12]. Therefore, a role for CEP78 in removal of CP110 from the centrosome is compatible with ultrastructural analyses in *Planarians*, which indicated that basal bodies fail to dock to the plasma membrane in CEP78 RNAi-depleted animals [34]. Although not addressed in detail in the current study, the elevated centrosomal CP110 levels observed in serum-fed CEP78 deficient cells are also compatible with previous work implicating CEP78 in regulation of PLK4-mediated centriole duplication [35].

We also uncovered a new physical interaction between CEP78 and the N-terminal region of CEP350, and found that CEP350 is important for recruitment of CEP78 to the centrosome as well as for its overall stability. The reduced interaction of the CEP78^L150S^ mutant protein with CEP350 might explain its lack of detection by western blot analysis of patient HSFs [30], since reduced interaction with CEP350 is likely to decrease the long-term stability of CEP78^L150S^. Our IP and IFM results furthermore suggests that CEP350 controls CP110 removal from the mother centriole not only via recruitment of TTBK2 [24], but also via CEP78-dependent recruitment of EDD1. Indeed, we observed reduced centrosomal levels of EDD1 in *CEP350* KO cells, consistent with this idea. It remains to be determined whether these two pathways act in sequence or in parallel to control centrosomal CP110 levels. Furthermore, as the recruitment of CEP350 to centrioles itself depends on FOP [24, 26, 38] and on the distal centriole protein C2CD3 [14], loss of these proteins may similarly affect the centrosomal recruitment of CEP78 and EDD1, and thereby negatively regulate CP110 levels via this pathway. It will be important to investigate these possibilities in future studies.

While increased CP110 levels can account for the decreased ciliation frequency observed in serum-deprived CEP78 mutant cells, it remains unclear why some of these cells form abnormally long cilia. Our analysis indicated that the ciliated mutant cells are able to locally displace or degrade CP110 at the mother centriole, but the underlying mechanism remains to be determined. Of note, the *CEP350* KO cells used in our study also seem to display a similar dual phenotype, since approximately 30% of these cells form cilia despite an inability to recruit TTBK2 and remove CP110 from the mother centriole [24]. Moreover, a recent genome-wide siRNA knockdown analysis showed that depletion of DDB1 caused an elongated cilia phenotype whereas depletion of EDD1 (UBR5) lead to fewer but normal length cilia [48], even though DDB1 and EDD1 are part of the same E3 ubiquitin ligase complex [49]. Thus, a slight imbalance in the regulation of this complex might determine whether or not cilia are formed and how long they are. It should also be noted that some studies have reported dual roles for CP110 during ciliogenesis, i.e. CP110 may not only function to inhibit ciliogenesis but may also promote ciliary extension in some contexts [44, 45]. Although the mechanism responsible for the elongated cilia phenotype seen in some of the CEP78 mutant cells remains unclear, one attractive candidate to be involved is CEP350, which we found to be present at abnormally high levels at the centrosome of ciliated *CEP78* KO cells, specifically. Consistent with this idea, we have observed that the CEP78 truncation mutant (clone #44), which expresses fewer, but normal length cilia, displays normal or slightly reduced levels of CEP350 at the centrosome in ciliated as well as in non-ciliated cells (data not shown). Alternatively, one or more substrates of EDD1 other than CP110 could also be affected in the *CEP78* mutant cells thereby affecting ciliary length control. For example, EDD1 was also shown to mediate ubiquitination of CSPP-L in turn promoting its recruitment to centriolar satellites and the centrosome [42]. CSPP-L is a well-known positive regulator of primary cilia formation [50] that was recently shown to bind directly to CEP104 to control axoneme extension [51]. Furthermore, CEP104 is implicated in ciliogenesis through interaction with the CEP97-CP110 complex and modulation of microtubule dynamics at the ciliary tip [51–53]. It will therefore be interesting to investigate if and how CSPP-L and CEP104 localization is affected in CEP78 mutant cells. Moreover, as CEP350 additionally binds to CYLD [54], a deubiquitinase that controls centriolar satellite homeostasis and ciliogenesis through de-ubiquitination of the E3 ubiquitin ligase MIB1 [55, 56], and was proposed to regulate axonemal IFT particle injection through FOP-CEP19-RABL2 interactions [24–26], it is tempting to speculate that CEP78 might impinge on these processes.

## Materials and methods

### PCR and cloning

For the establishment of the N-terminal FLAG-tagged CEP78 construct a cDNA clone coding for human full-length CEP78 cDNA (GeneCopoiea pShuttle™Gateway® PLUS ORF, NCBI accession number NM_001098802) was used as template for amplification with the Phusion High-Fidelity DNA polymerase (New England BioLabs) under standard PCR conditions and attB forward and reverse primers (see Key Resources Table) compatible with the Gateway cloning technology (Invitrogen/Thermo Fisher Scientific) for inclusion of the insert into the pDonor201 Gateway vector (Invitrogen/Thermo Fisher Scientific). To produce human FLAG-CEP78^L150S^, site directed mutagenesis was performed with QuickChange Lightning Site-Directed Mutagenesis Kit (Agilent) and primers CEP78 mut_F and CEP78 mut_R (see Key Resources Table). Accuracy of the *CEP78* ORF was assessed by Sanger sequencing. Subsequently, the desired fragment was cloned into p3xFLAG-CMV/DEST (gift from Dr. Ronald Roepman lab, Radboud University Medical School, Nijmegen, NL; see Key Resources Table for details) to produce p3xFLAG-CEP78 or p3xFLAG-CEP78^L150S^.

Plasmids encoding full-length and truncated versions (N-terminus, middle, N-terminus plus middle, C-terminus) of CEP78 tagged with GFP in the N-terminus were generated by PCR using relevant primers (see Key Resources Table) and pFLAG-CEP78 as template; PCR products were cloned into pEGFP-C1 by standard approaches, following digestion with BamHI (Roche, cat# 1056704001) and KpnI (Roche, cat#10899186001) and ligation with T4 DNA ligase (Applichem, cat# A5188). Ligated plasmids were transformed into competent *E. coli* DH10B cells and cells harboring recombinant plasmids selected on Luria Bertani (LB; Sigma-Aldrich) agar plates with 50 μg/ml kanamycin. Plasmids were purified using Plasmid DNA Mini Kit I (Omega Biotech, cat# D6943-02) or Nucleobond Xtra Mid kit (Macherey-Nagel, cat# 740410.50), according to protocols supplied by the manufacturers. Sequences of plasmid inserts were verified by Eurofins Genomics.

Gateway system compatible pENTR vectors encoding for mNeonGreen-CEP78 or mNeonGreen-CEP78L150S fusions were generated by subcloning of ORFs from pEGFP constructs described above in frame with mNeonGreen using the BamH1/Kpn1 sites. For lentivirus particle production these plasmids were recombined with pCDH-EF1a-Gateway-IRES-BLAST plasmids as described in [51] using LR Clonase II (Invitrogen/Thermo Fisher Scientific).

### Mammalian cell culture and transfection

Human skin fibroblasts were cultured as described previously [29, 30]. Human embryonic kidney-derived 293T cells were cultured and transfected with plasmids as described previously [61]. RPE1 cells were grown in T-75 flasks in a 95% humidified incubator at 37 °C with 5% CO2. Cells were cultured in Dulbecco’s Modified Eagle Medium (DMEM) (GIBCO®, cat# 41966-029) supplemented with 10% fetal bovine serum (FBS) and 1% penicillin/streptomycin (P/S). When cells were about 80-90% confluent, they were passaged and set up for new experiments. Upon seeding, RPE1 cells were washed once with preheated 1 x phosphate buffered saline (PBS) and then incubated with 1% Trypsin-EDTA (ethylenediaminetetraacetic acid) (Sigma-Aldrich, cat# T4174) solution for 5 min at 37 °C. The detached cells were aspired in an appropriate volume of new preheated DMEM and seeded into petri dishes with or without sterile glass coverslips [62]. When the cells had reached 80-90% confluency they were serum starved for 24 hours to induce formation of primary cilia. A small amount of cell solution was passaged on in a new T-75 flask containing new preheated DMEM. All cell lines were tested negative for Mycoplasma and those not generated in this study have been used in prior publications. New cell lines generated in this study were validated by sanger sequencing.

For transfection with plasmids, RPE1 cells were seeded in 60 mm petri dishes as described above to about 50-70% confluency at the time of transfection. For rescue experiments, 2 μg pFLAG-CEP78 or pFLAG-CEP78^L150S^ and 6 μl FuGENE®6 were mixed in 100 μl serum- and antibiotic free DMEM and incubated for 20 minutes at room temperature. New DMEM was added to the cells prior to transfer of the transfection reagent, which was gently dripped onto the cells and swirled to make sure it was dispersed well. After cells were incubated with transfection medium for 4 hours, the medium was replaced with serum free DMEM for 24 hours to induce growth arrest. Gene silencing in RPE1 cells by siRNA was performed using DharmaFect (see Key Resources Table). Briefly, RPE1 cells were seeded in DMEM medium with serum and antibiotics in 30 mm petri dishes at 20-25% confluency. Transfection was performed by combining 50 nM siRNA with 5 µl of DharmaFect in 200 µl of serum and antibiotic-free DMEM medium for 15 min at room temperature. After incubation, the complexes were added dropwise onto the cells and the media was swirled gently to ensure good dispersion prior to incubation at 37 °C. Cells were kept under these conditions for 48 hours and then serum starved for 24 hours before being processed for further experiments.

RPE1 cell transduction with lentivirus particles and Blasticidine selection (10µg/ml f.c.) was conducted as described in [51].

### Immunofluorescence microscopy and image analysis

Standard epifluorescence IFM analysis was performed as described previously [62]. Briefly, glass coverslips containing cells of interest were washed in PBS and fixed with either 4% paraformaldehyde (PFA) solution for 15 min at room temperature or on ice, or with ice-cold methanol for 10-12 min at -20 °C. After three washes in PBS the PFA-fixed cells were permeabilized 10 min in 0.2% Triton X-100 and 1% Bovine Serum Albumin (BSA) in PBS and blocked in 2% BSA in PBS for 1 hour at room temperature. Coverslips were incubated overnight at 4 °C in primary antibodies (see Key Resources Table) diluted in 2% BSA in PBS. The next day, coverslips were washed three times with PBS for 5 min and incubated for 1 hour at room temperature in secondary antibodies (see Key Resources Table) diluted in 2% BSA in PBS. Coverslips were washed three times in PBS for 5 min before staining with 2 μg/ml DAPI solution PBS for 30 sec. Finally, coverslips were washed once with PBS and before mounting on objective glass with 6% n-propyl gallate diluted in glycerol and 10x PBS and combined with Shandon^TM^ Immuno-Mount^TM^ (Thermo Scientific, cat# 9990402) in a 1:12 ratio. Cells were imaged on a motorized Olympus BX63 upright microscope equipped with a DP72 color, 12.8 megapixel, 4140 x 3096 resolution camera. cellSense Dimension software 1.18 from Olympus was used to measure cilia length; ImageJ version 2.0.0-rc-69/1.52i was used to measure the relative mean fluorescence intensity (MFI) of relevant antibody-labeled antigens at the centrosome/basal body. A fixed circle was drawn around a centrosome in a cell. This same region of interest (ROI) was used to measure the MFI of a specific protein at the centrosome. A constant ROI was also drawn to measure the cell background signal which was subtracted from the MFI measured at the centrosome/basal body. Images were prepared for publication using Adobe Photoshop® and Adobe Illustrator®.

Super resolution imaging of RPE1 cells was conducted as described in [51] except that cells were mounted in ProLong Diamond Antifade mountant. Settings for hardware-assisted axial and software-assisted lateral channel alignment and image reconstruction were validated by imaging of RPE1 cells stained for CEP164 and labeled with secondary antibodies conjugated to either Alexa488, DyeLight550, or Alexa647. For superimposing 3D-SIM images on electron microscopy micrographs of centrioles (kindly provided by Dr. Michel Bornens; [40]), 3D-SIM images were sized to same digital pixel resolution as original EM images using the bicubic algorithm in ImageJ and maximal intensity z-projections of single channel images overlaid on micrographs using centriole centers / centriole distal ends as reference points.

### Statistical analysis

Statistical analyses were performed using GraphPad Prism 6.0. The background-corrected MFI measured at the centrosome was normalized to relevant control cells. Mean and standard deviation (SD) was calculated for all groups, and outliers were identified and removed by the ROUT method before the statistical tests were conducted. The data was tested for gaussian normality using either D’Agostino’s K-squared test or Shapiro-Wilk test. Depending on the distribution of the data, two-tailed and unpaired Student’s t-test or Mann Whitney test were used when comparing two groups. Also, depending on the distribution of the data, One-way ANOVA followed by Tukey’s, Dunnet’s or Dunn’s multiple comparison tests was used when comparing more than two groups. Unless otherwise stated, the statistical analyses were performed on at least three independent biological replicates (n=3). A p-value under 0.05 was considered statistically significant and p-values are indicated in the figures with asterisks as follows; * p<0.05, ** <p0.01, ***<0.001, and **** p<0.0001.

### Immunoprecipitation (IP), protein quantification, SDS-PAGE and western blot analysis

Immunoprecipitation of transfected 293T cell lysates was performed as described previously [61] using relevant antibody-conjugated beads (see Key Resources Table). Protein concentrations were measured using the *DC*^TM^ Protein Assay Kit I from Bio-Rad (cat # 5000111) by following the manufacturer’s protocol. For SDS-PAGE analysis of RPE1 and HSF cells, protein samples were prepared by lysis of cells with 95 °C SDS-lysis buffer (1% SDS, 10 mM Tris-HCl, pH 7.4) and cell lysates transferred to Eppendorf tubes and heated shortly at 95 °C. The samples were then sonicated two times for 30 seconds to shear DNA followed by centrifugation at 20,000 x *g* for 15 min at room temperature to pellet cell debris. Supernatants were transferred to new Eppendorf tubes and an aliquot of each sample used for determination of protein concentration. Samples with equal concentrations of protein were prepared for SDS-PAGE analysis by addition of NuPAGE^TM^ LDS Sample Buffer (4X) from Thermo Fisher Scientific (cat# NP0007) and 50 mM DTT. Samples from IP analysis in 293T cells, performed using modified EBC buffer [61], were prepared for SDS-PAGE in a similar fashion. Protein samples were heated at 95 °C for 5 minutes before loading them on a Mini-PROTEAN® TGX^TM^ Precast Gel 4-15% from Bio-Rab Laboratories, Inc. (cat# 456-1083 or cat #456-1086). PageRuler^TM^ Plus Prestained Protein Ladder (Thermo Fisher Scientific, cat# 26619) was used as molecular mass marker. SDS-PAGE was performed using the Mini-PROTEAN® Tetra System from Bio-Rad. Gels were run at 100 V for 15 minutes and 200 V for 45 minutes. Using the Trans-BLOT® Turbo Transfer System from Bio-Rad Laboratories, Inc. the proteins were transferred at 1.3 A, 25 V for 10 min from the gel to a Trans-Blot® Turbo^TM^ Transfer Pack, Mini format 0.2 μm Nitrocellulose membrane (cat# 1704158). Ponceau-S solution was added to the membrane to visualize the proteins before blocking in 5% milk in Tris-buffered saline with Tween-20 (TBS-T; 10 mM Tris-HCl, pH 7.5, 150 mM NaCl, 0.1% Tween-20) for 2 hours at room temperature. Primary antibodies (see Key Resources Table) were diluted in 5% milk in TBS-T and incubated with the membrane overnight at 4°C. The membrane was washed three times 10 min at room temperature in TBS-T on a shaker before incubation with secondary antibody (see Key Resources Table), diluted in 5% milk in TBS-T, for 1 hour at room temperature. The membrane was washed three times 10 min at room temperature in TBS-T on a shaker. Finally, SuperSignal^TM^ west Pico PLUS chemi-luminescent Substrate (Thermo Scientific, cat # 34580) was mixed in a 1:1 ratio and added to the membrane for 5 min before development on a FUSION FX SPECTRA machine from Vilber.

### Quantitative mass spectrometry (MS) and peptide identification

For SILAC-based mass spectrometry, three distinct SILAC culture media were used: light- (Lys0 and Arg0); medium- (Lys4 and Arg6) and high-labeled media (Lys8 and Arg10). 293T cells were seeded in 100 mm petri dishes and SILAC labeled for 1 week to ensure proper amino acid incorporation. After the incorporation phase, the cells were transfected with 2 µg of plasmid encoding FLAG-Ap80 (control), FLAG-CEP78 or FLAG-CEP78^L150S^ using 6 μl FuGENE®6, as described above, and then subjected to lysis using modified EBC buffer [56] and FLAG IP as described above. Affinity enriched proteins were digested with trypsin and the resulting peptides desalted prior to LCMS analysis using an Easy-nLC 1000 system (Thermo Scientific) connected to a Q Exactive HF-X mass spectrometer (Thermo Scientific). Peptides were separated by a 70 minutes gradient using increased buffer B (95% ACN, 0.5% acetic acid). The instrument was running in positive ion mode with MS resolution set at 60,000 for a scan range of 300 to 1700 m/z. Precursors were fragmented by higher-energy collisional dissociation (HCD) with normalized collisional energy of 28 eV. For protein identification and quantitation, the obtained MS raw files were processed by MaxQuant software version 1.6.1.0 [60] and searched against a FASTA file from UniProt.

### RNAseq analysis

RNA was extracted using the Direct-zol RNA Miniprep Kit (Zymo Research) following the manufacturer’s instructions. RNA degradation and contamination were monitored on 1% agarose gels. RNA purity was checked using the NanoPhotometer® spectrophotometer (IMPLEN, CA, USA). RNA concentration was measured using Invitrogen^TM^ Qubit® RNA Assay Kit in Qubit® 2.0 Fluorometer (Thermo Fisher Scientific). RNA integrity was assessed using the Agilent RNA Nano 6000 Assay Kit of the Bioanalyzer 2100 system (Agilent, CA, USA). A total amount of 3 µg RNA per sample was used as input material for the RNA sample preparations. Sequencing libraries were generated using NEBNext® Ultra™ RNA Library Prep Kit for Illumina® (New England BioLabs, USA) following manufacturer’s recommendations and index codes were added to attribute sequences to each sample. Briefly, mRNA was purified from total RNA using poly-T oligo-attached magnetic beads. Fragmentation was carried out using divalent cations under elevated temperature in NEBNext First Strand Synthesis Reaction Buffer (5X). First strand cDNA was synthesized using random hexamer primer and M-MuLV Reverse Transcriptase (RNase H -). Second strand cDNA synthesis was subsequently performed using DNA Polymerase I and RNase H. Remaining over-hangs were converted into blunt ends via exonuclease/polymerase activities. After adenylation of 3’ ends of DNA fragments, NEBNext Adaptor with hairpin loop structure were ligated to prepare for hybridization. In order to select cDNA fragments of preferentially 150∼200 bp in length, the library fragments were purified with AMPure XP system (Beckman Coulter Life Sciences). Then 3 µl USER® Enzyme (New England BioLabs) was used with size-selected, adaptor-ligated cDNA at 37°C for 15 min followed by 5 min at 95 °C before PCR. Then PCR was performed with Phusion® High-Fidelity DNA polymerase, Universal PCR primers and Index (X) Primer (New England BioLabs). At last, PCR products were purified (AMPure XP system) and library quality was assessed on the Agilent Bioanalyzer 2100 system. The clustering of the index-coded samples was performed on a cBot Cluster Generation System using HiSeq PE Cluster Kit cBot-HS (Illumina) according to the manufacturer’s instructions. After cluster generation, the library preparations were sequenced on an Illumina Hiseq platform and 125 bp/150 bp paired-end reads were generated. Raw data (raw reads) of fastq format were firstly processed through in-house perl scripts. In this step, clean data (clean reads) were obtained by removing reads containing adapter, reads containing poly-N and low quality reads from raw data. At the same time, Q20, Q30 and GC content the clean data were calculated. All the downstream analyses were based on the clean data with high quality. Reference genome and gene model annotation files were downloaded from genome website directly. Index of the reference genome was built using Bowtie v2.2.3 [63] and paired-end clean reads were aligned to the reference genome using TopHat v2.0.12 [64]. We selected TopHat as the mapping tool for that TopHat can generate a database of splice junctions based on the gene model annotation file and thus a better mapping result than other non-splice mapping tools. HTSeq v0.6.1 [65] was used to count the reads numbers mapped to each gene and then Fragments Per Kilobase of transcript sequence per Millions base pairs sequenced (FPKM) of each gene was calculated based on the length of the gene and reads count mapped to this gene. FPKM considers the effect of sequencing depth and gene length for the reads count at the same time, and is currently the most commonly used method for estimating gene expression levels [66]. Prior to differential gene expression analysis, for each sequenced library, the read counts were adjusted by edgeR program package through one scaling normalized factor [67]. Differential expression analysis of two conditions was performed using the DEGSeq R package (1.20.0) [68].

## Acknowledgements

We thank Søren Lek Johansen and Maria Schrøder Holm for expert technical assistance, Benedicte Schultz Kappel and Frida Roikjer Rasmussen for help with plasmid generation, and Nynne Christensen, Center for Advanced Bioimaging (CAB) Denmark, for help with 3D-SIM acquisition. Dr. Kay Schink kindly provided plasmids for lentivirus particle generation and assisted in 3D-SIM acquisition at OUH Radiumhospitalet. We are grateful to Drs. Peter K. Jackson, Tomoharu Kanie, Francesc Garcia-Gonzalo, Laurence Pelletier, Brian David Dynlacht, Anne-Marie Tassin, Maria Gavilan, Rosa M. Rios, Éric A. Cohen and Peter ten Dijke for reagents, and Dr. Michel Bornens for sharing original EM microcraphs. This study was supported grants from Independent Research Fund Denmark (#8020-00162B to PF and LBP, and #8021-00425A to JSA), from the Carlsberg Foundation (#CF18-0294 to LBP) and from the Swiss National Science Foundation (#176097 to CR).

## Supplementary figure legends

**Figure 1-figure supplement 1.** Western blot analysis and ciliary frequency and length in RPE1 WT and *CEP78* KO clones. **(A)** Western blot analysis of serum-deprived RPE1 WT and four different *CEP78* KO clones, generated by CRISPR/Cas9 methodology. Blots were probed with antibodies against CEP78 and GAPDH (loading control), as indicated. The lower band in the CEP78 blot is due to unspecific staining of the antibody. Molecular mass markers are indicated in kDa to the right. Unless otherwise indicated, clone #73 was used for all experiments in RPE1 cells performed in this manuscript, and is designated *CEP78* KO. (**B, C**) Quantification of ciliary frequency (B) and length (C) in serum-deprived RPE1 WT and different *CEP78* KO clones. Differences in cilia numbers and size between the WT and the several *CEP78* KO clones were addressed through accumulated data from three independent experiments (for cilia frequency: n= 304 for WT; n=308 for clone #2; n=321 for clone #44 and n=311 for clone #52; and for ciliary length: n=145 for WT; n=68 for clone #2; n=98 for clone #44 and n=62 for clone #52) by performing an ordinary one-way ANOVA with Dunnet’s multiple comparison test amongst all groups, in relation to WT cells mean (control group). Error bars in (B, C) indicate mean ± SD. (*) p<0.05; (**) p<0.01; (***) p<0.001.

**Figure 2-figure supplement 1.** Expression of FLAG-CEP78 and FLAG-CEP78^L150S^ in RPE1 WT and *CEP78 KO* cells. **(A)** FLAG blots from serum-deprived WT and *CEP78* KO lysates transfected with indicated FLAG fusions. GAPH was used as a loading control. (**B**) Quantification of data depicted in (A). Graphs of the FLAG blots are presented as the mean + SD from three individual experiments analyzed in duplicates. Ordinary one-way ANOVA with Tukey’s multiple comparison test was used for the statistical analysis, where no significant statistical differences were reported between the groups.

**Figure 2-figure supplement 2.** CEP78 endogenous antibody binds equally well to FLAG-CEP78 WT and L150S mutant fusions. 293T cells transiently expressing FLAG-CEP78 or FLAG-CEP78^L150S^ were subjected to IP with FLAG. Pellet fractions were analyzed by SDS-PAGE and western blotting using antibodies against FLAG and endogenous CEP78. Molecular mass marker is indicated in kDa to the left of the blots.

**Figure 2-figure supplement 3.** Stable expression of mNG-CEP78 and mNG-CEP78^L150S^ in RPE1 WT and *CEP78* KO cells. **(A)** Western blots of lysates of serum-deprived WT and *CEP78* KO RPE1 cells lines stably expressing the indicated mNG-tagged fusions. The blots were probed with antibodies against CEP78 (upper panel) and *α*-tubulin (loading control; lower panel). Molecular mass markers are shown in kDa to the left. (**B**) Representative light (upper panels) and fluorescence (lower panels) images of live cells expressing the indicated fusions. Note that mNG-CEP78 is concentrated at the centrosome whereas mNG-CEP78^L150S^ is not. Scale bar, 20 μm.

**Figure 3-figure supplement 1.** CEP78 associates independently with CEP350 and VPRBP. (**A**) 293T cells expressing the indicated Myc- or GFP-fusions were subjected to IP with Myc antibody-coated beads and input and pellet (IP Myc) fractions analyzed by SDS-PAGE and western blotting with VPRBP, Myc, FOP or GFP antibodies as indicated. (**B**) Similar analysis as in (A), but using GFP antibody-coated beads for IP instead of Myc beads. Molecular mass markers are indicated in kDa to the left of the blots. Arrowhead in (A) indicates IgG band observed in the lower part of the GFP blot.

**Figure 4-figure supplement 1.** Validation of *CEP350* KO cells by IFM analysis. (**A**) RPE1 control and *CEP350* KO cells were serum starved for 24 hours before fixation and staining with the indicated antibodies and DAPI for nuclear visualization. Insets show closeups of the centrosomal/ciliary area (dashed boxes). Scale bars: 5 µm in representative images and 1 µm in closeups. (**B**) Quantification of CEP350 mean fluorescence intensity (MFI) at the centrosome, based on images as shown in (A). A total of 38 and 26 cells were analyzed for control and *CEP350* KO cells, respectively. For statistical analysis, the data was first normalized to the control group values and differences were assessed using an unpaired and two-tailed Student’s t-test. Data is shown as the mean ± SD. a.u., arbitrary units; (**) p<0.01.

**Figure 6-figure supplement 1.** VPRBP centrosomal levels are increased or unchanged in RPE1 *CEP78* KO cells. (**A, C**) Representative IFM micrographs of ciliated (**A**) and non-ciliated (**C**) serum-starved RPE1 WT and *CEP78* KO cells labeled with the antibodies indicated in the figure. DAPI was used to mark the nucleus (blue). Insets show enlarged views of the cilium-centrosome region. Scale bars: 5 μm in original images and 1 μm in closeups. (**B, D**) Quantification of the VPRBP MFI at the centrosome based on images shown in (A) and (C) based on data from three independent experiments (n= 156 and n=158 for ciliated WT and CEP78 KO cells, respectively; n= 162 and n= 166 for non-ciliated WT and CEP78 KO cells, respectively) using a two-tailed and unpaired Student’s t-test. Data is shown as mean ± SD and was first normalized in relation to WT values. a.u., arbitrary units; (**) p<0.01; n.s., not statistically significant.

**Figure 6-figure supplement 2.** EDD1 centrosomal levels are reduced in RPE1 *CEP350* KO cells. (**A, C**) Representative IFM micrographs of ciliated (**A**) and non-ciliated (**C**) serum-starved RPE1 control and *CEP350* KO cells labeled with the antibodies indicated in the figure. DAPI was used to mark the nucleus (blue). Insets show enlarged views of the cilium-centrosome region. Scale bars: 5 μm in original images and 1 μm in closeups. (**B, D**) Quantification of the EDD1 MFI at the centrosome based on images as shown in (A) and (C) using a two-tailed and unpaired Student’s t-test from two independent experiments (n=105 and n= 111 for ciliated WT and *CEP78* KO cells, respectively; n=118 for non-ciliated WT and CEP78 KO cells). Data is shown as mean ± SD and was first normalized in relation to WT values. a.u., arbitrary units; (****) p<0.0001.

**Figure 7-figure supplement 1.** Elevated cellular CP110 levels in RPE1 *CEP78* KO clones. **(A)** Western blot analysis of serum deprived RPE1 WT and three different *CEP78* KO clones, as indicated. Blots were probed with antibodies against CP110 and GAPDH (loading control). Molecular mass markers are indicated in kDa to the right. (**B**) Quantification of relative cellular CP110 levels based on blots as shown in (A) from two independent experiments. The means of the CEP78 KO clones, were compared in relation to the WT mean, selected as the control group using an ordinary one-way ANOVA followed by a Dunnet’s multiple comparison test. Error bars of graphs represent SD and data are shown as mean ± SD. (**C**) Representative western blot of WT and *CEP78* KO cells (KO; clone #73) stably expressing mNG-CEP78 or mNG-CEP78^L150S^ as indicated. Blots were probed with indicated antibodies (*α*-tub=*α*-tubulin) and the molecular mass markers are indicated in kDa to the right of the blot. Quantification of relative CP110 band intensities revealed no significant changes between the three conditions; CP110 level relative to that of WT mNG-CEP78 was 95% +/-12% for the *CEP78* KO mNG-CEP78 line and 90% +/- 9% for the *CEP78* KO mNG-CEP78^L150S^ line, respectively (n=3). Blots were probed with indicated antibodies.

**Figure 7-figure supplement 2.** Serum-fed CEP78 deficient cells display elevated cellular and centrosomal levels of CP110. (**A**) CP110 western blot analysis of lysates from RPE1 WT and *CEP78* KO cells grown in the presence of serum; α-tubulin serves as loading control. (**B**) Quantification of the CP110 blots depicted in (A) based on four individual experiments analyzed in duplicates. Error bars indicate SD. Student’s t-test (unpaired, two-tailed) was used to analyze the differences between both cellular groups. (**C**) Representative IFM images of cycling RPE1 WT and *CEP78* KO cells labeled with the indicated antibodies. DAPI was used to mark the nucleus. Insets show closeup images of the centrosome region. Scale bars: 5 µm in original images and 1 µm in closeups. (**D**) Data quantification of the relative MFI of CP110 based on images as shown in (C). Data is present as mean ±SD. Statistical analysis was performed using Student’s t-test (unpaired, two-tailed) of three independent experiments (n=158 for WT and n= 147 for *CEP78* KO cells). a.u., arbitrary units; (*) p<0.05; (****) p<0.0001.

**Figure 7-figure supplement 3.** Barplots showing expression of *CP110, CEP350, VRPBP* and *EDD1* genes for patients (n=3) and controls (n=4), based on RNA-seq data of skin fibroblasts. Expression levels are shown in fpkm (Fragments Per Kilobase of transcript sequence per Millions base pairs sequenced). The patient data are combined data from HSFs derived from three different *CEP78* probands; F3 individual II:1 described in [30] and KN10 individual II-1 and 2716s15 individual II-2 described in [29]. Control data are combined data from HSFs from four unaffected individuals described in the same studies. Nominal p-values derived from unpaired two-tailed t-tests are shown on top of the bars. None of the comparisons were significantly different (p<0.05).

**Figure 7-figure supplement 4.** Relative CP110 levels in WT and *CEP78* KO cells treated with cycloheximide. (**A**) Representative western blot of lysates from serum-deprived WT and *CEP78* KO cells treated with 200 ng/ml cycloheximide (CHX) for the indicated times. The drug was added at the onset of serum deprivation (time 0 h). Blots were probed with CP110 and GAPDH (loading control) antibodies; molecular mass markers are shown in kDa to the left. (**B**) Quantification of average CP110 levels determined based on western blots as shown in (A). The analysis is based on data from four independent experiments, normalized to the relative CP110 level for each cell type at time zero. Note that the data indicates decline in CP110 levels with time in the WT cells but not in the *CEP78* KO cells; however, due to fluctuations from experiment to experiment the changes were not statistically significant (ns; p values denoted in parenthesis above the graphs). The data is presented as mean ±SD.

**Figure 8-figure supplement 1.** Validation of CP110 knockdown in RPE1 WT and *CEP78* KO cells by IFM. (A, B) Quantification of CP110 centrosomal levels in ciliated and non-ciliated WT and *CEP78* KO cells treated with control (siMock) or CP110-specific siRNA (siCP110). Both graphs are based on one biological experiment with 35 observations of ciliated and 35 observations of non-ciliated cells per condition. Statistical analysis was performed using and Ordinary One-way ANOVA followed by a Dunnett’s multiple comparison test amongst all groups in relation to WT siMock, designated as the control group. Differences between *CEP78* KO cells treated with siMock or siCP110 were addressed by performing an unpaired and two-tailed Student’s t-test. Bars are represented as mean ±SD. (a.u.) arbitrary units; (*) p<0.05; (**) p<0.01.

## Additional files

**Figure 1-source data 1.** Original western blots corresponding to Figure 1G. The upper blots are labeled with pRb antibody and lower blots with GAPDH antibody.

**Figure 1-source data 2.** Original western blots corresponding to Figure 1J. The upper blots are labeled with pRb antibody and lower blots with GAPDH antibody.

**Figure 3-source data 1.** Raw data from the CEP78 interactome analysis depicted in Figure 3A. 293T cells grown in SILAC medium and expressing FLAG-CEP78 WT, FLAG-CEP78^L150S^ or FLAG-Ap80 (negative control) were subjected to FLAG IP and pellets analyzed by mass spectrometry.

**Figure 3-source data 2.** Original western blots corresponding to Figure 3B. Upper left, input Myc blot; upper middle, input 100 kDa FLAG blot; upper right, IP Myc and 100 kDa FLAG blots; bottom left, input 35 kDa FLAG blot; bottom right, IP 35 kDa FLAG blot.

**Figure 3-source data 3.** Original western blots corresponding to Figure 3C. Upper left, input Myc blot; upper middle, IP Myc blot; upper right, input GFP blot; bottom left, IP GFP blot; bottom middle, inpot FOP blot; bottom right, IP FOP blot.

**Figure 3-source data 4.** Original western blots corresponding to Figure 3D. Upper left, input Myc blot; upper right, IP Myc blot; lower left, input GFP blot; lower right, IP GFP blot.

**Figure 4-source data 1.** Original western blots for Figure 4B. Left, CEP78 blot; right, *α*-tubulin blot.

**Figure 6-source data 1.** Original western blots for Figure 6A. Left, VPRBP blot; right, *α*-tubulin blot.

**Figure 7-source data 1.** Original western blots for Figure 7A. Left, CP110 blot; right, GAPDH blot.

**Figure 8-source data 1.** Original bots for Figure 8A. Left, CP110 blot; right, *α*-tubulin blot. **Figure 1-figure supplement 1-source data 1.** Original western blots for Figure 1-figure supplement 1. Left, CEP78 blot; right, GAPDH blot.

**Figure 2-figure supplement 1-source data 1.** Original western blots for Figure 2-figure supplement 1. Left, FLAG blot; right, GAPDH blot.

**Figure 2-figure supplement 2-source data 1.** Original western blots for Figure 2-figure supplement 2. Left, FLAG blot; right, CEP78 blot.

**Figure 2-figure supplement 3-source data 1.** Original western blots for Figure 2-figure supplement 3. Left, CEP78 blot; right, *α*-tubulin blot

**Figure 3-figure supplement 1-source data 1.** Original western blots for Figure 3-figure supplement 1A. Top row from left to right: input VPRBP blot, IP VPRBP blot, input Myc blot. Second row from left to right: IP Myc blot, input FOP blot, IP FOP blot. Third row from left to right: input GFP blot (upper), IP GFP blot (upper). Fourth row left to right: input GFP blot (lower), IP GFP blot (lower).

**Figure 3-figure supplement 1-source data 2.** Original western blots for Figure 3-figure supplement 1B. Top row, input VPRBP blot; second row, IP VPRBP blot; third row left, input Myc blot; third row right, IP Myc blot; fourth row left, input GFP blots; fourth row right, IP GFP blots.

**Figure 7-figure supplement 1-source data 1.** Original western blots for Figure 7-figure supplement 1. Left, CP110 (upper) and GAPDH blots in panel A. Right, CP110 (upper) and *α−*tubulin (lower) blots in panel C.

**Figure 7-figure supplement 2-source data 1.** Original western blots for Figure 7-figure supplement 2A. Top, CP110 blot; bottom, *α−*tubulin blot.

**Figure 7-figure supplement 3-source data 1.** Raw RNA seq data for Figure 7-figure supplement 3.

**Figure 7-figure supplement 4-source data 1.** Original western blots for Figure 7-figure supplement 4A. Upper left, WT CP110 blot; upper right, WT GAPDH blot. Lower left, *CEP78* KO CP110 blot; lower right, *CEP78* KO GAPDH blot.

## Appendix

### Key Resources Table

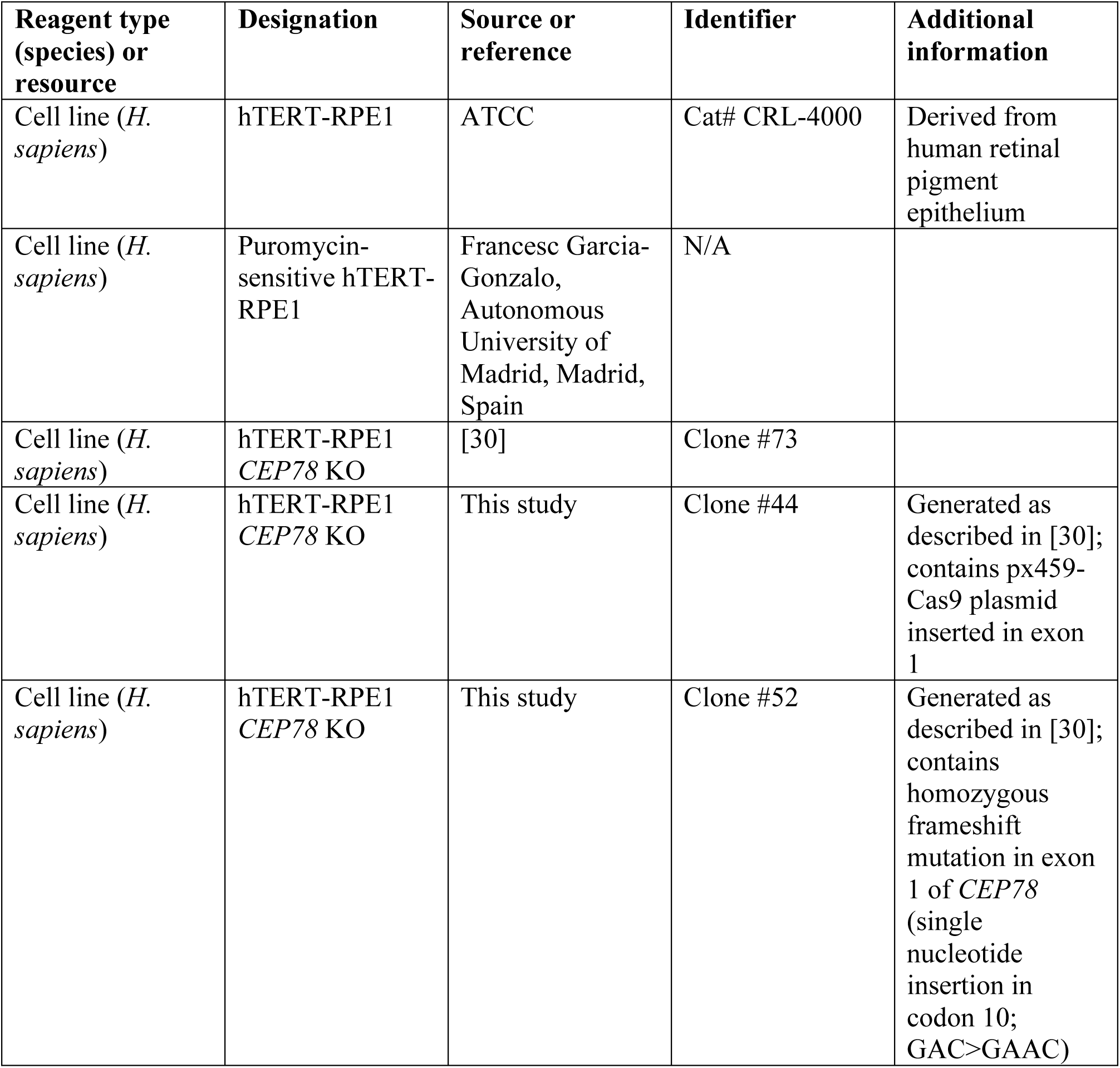
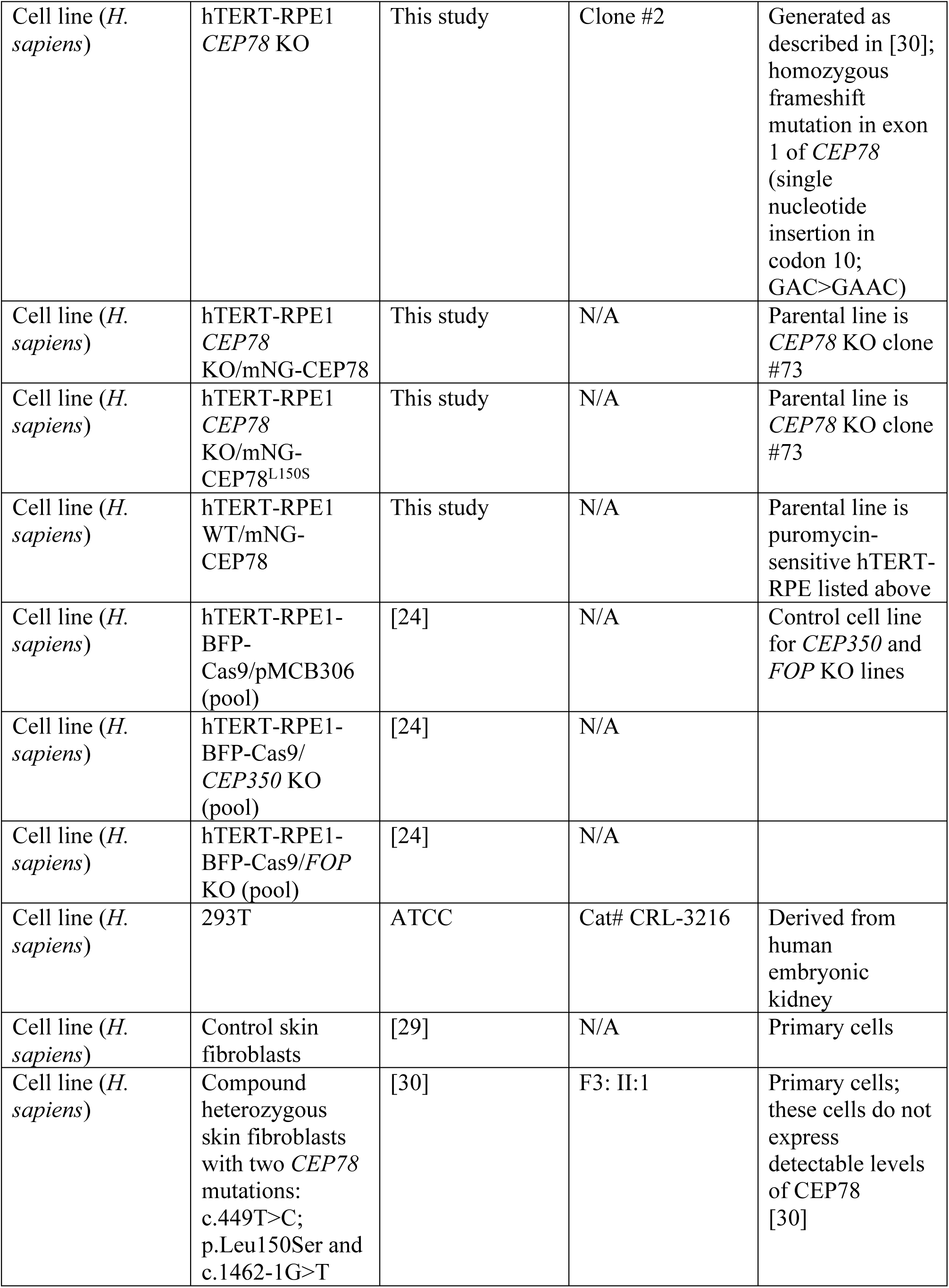
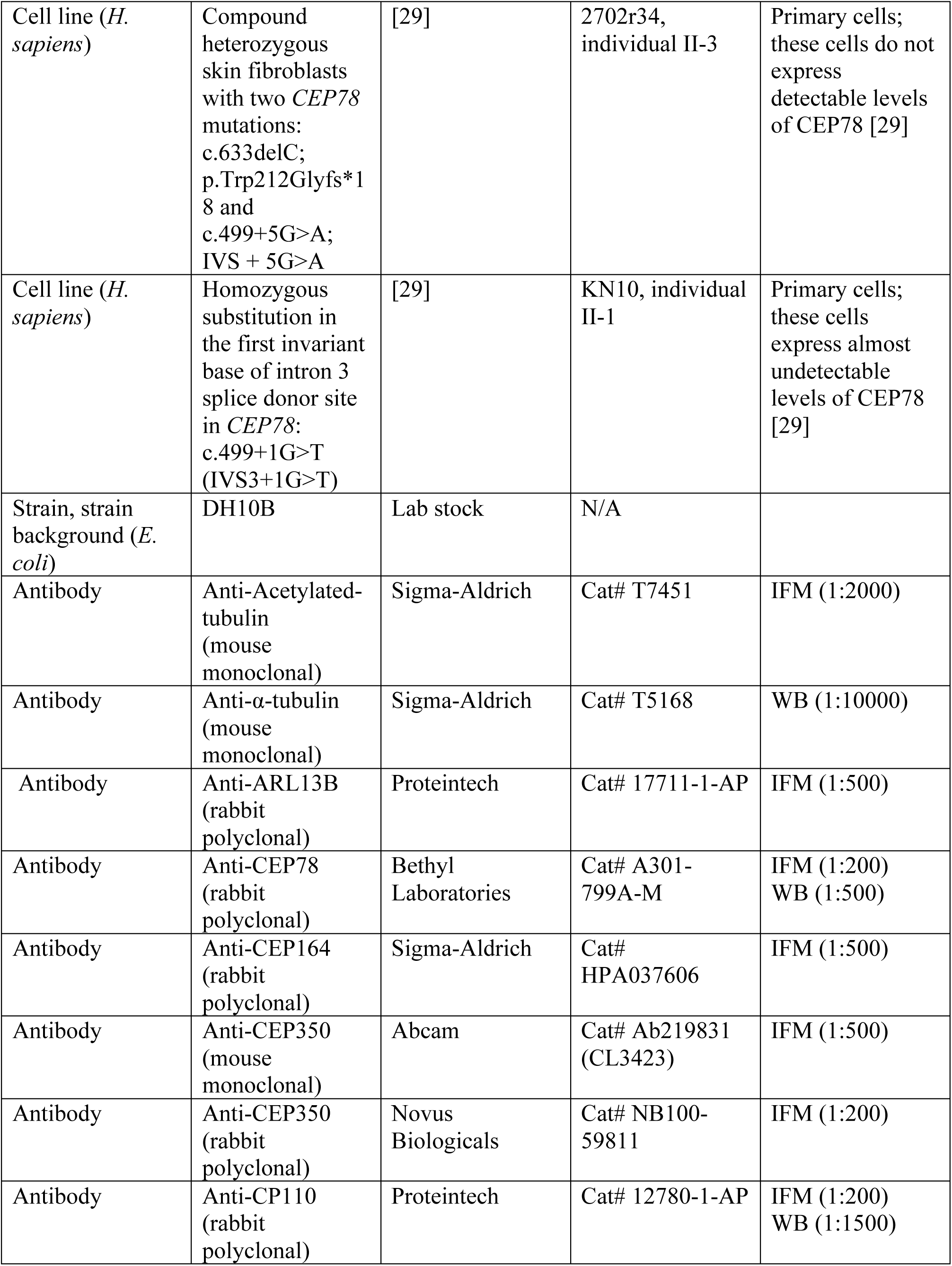
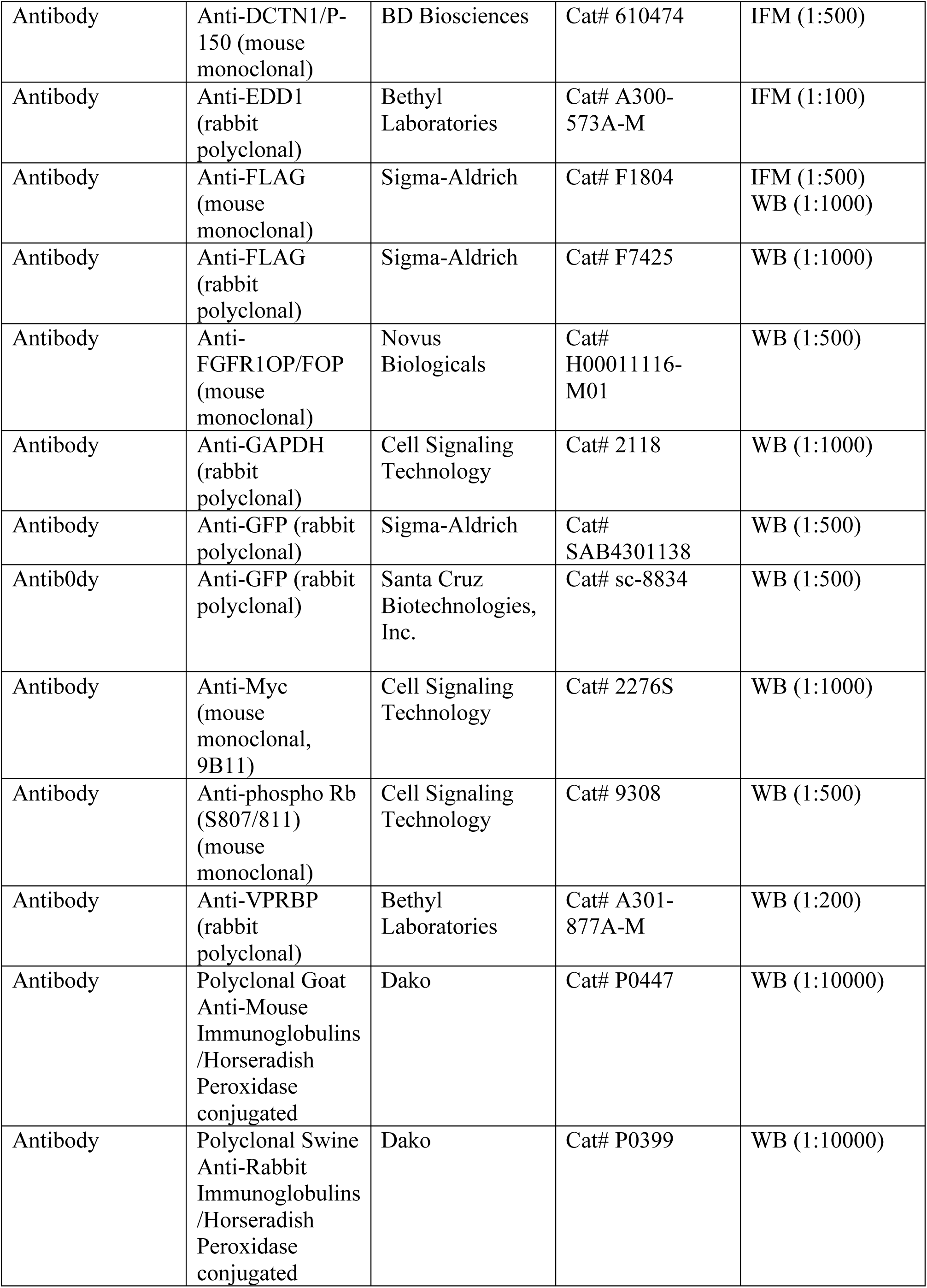
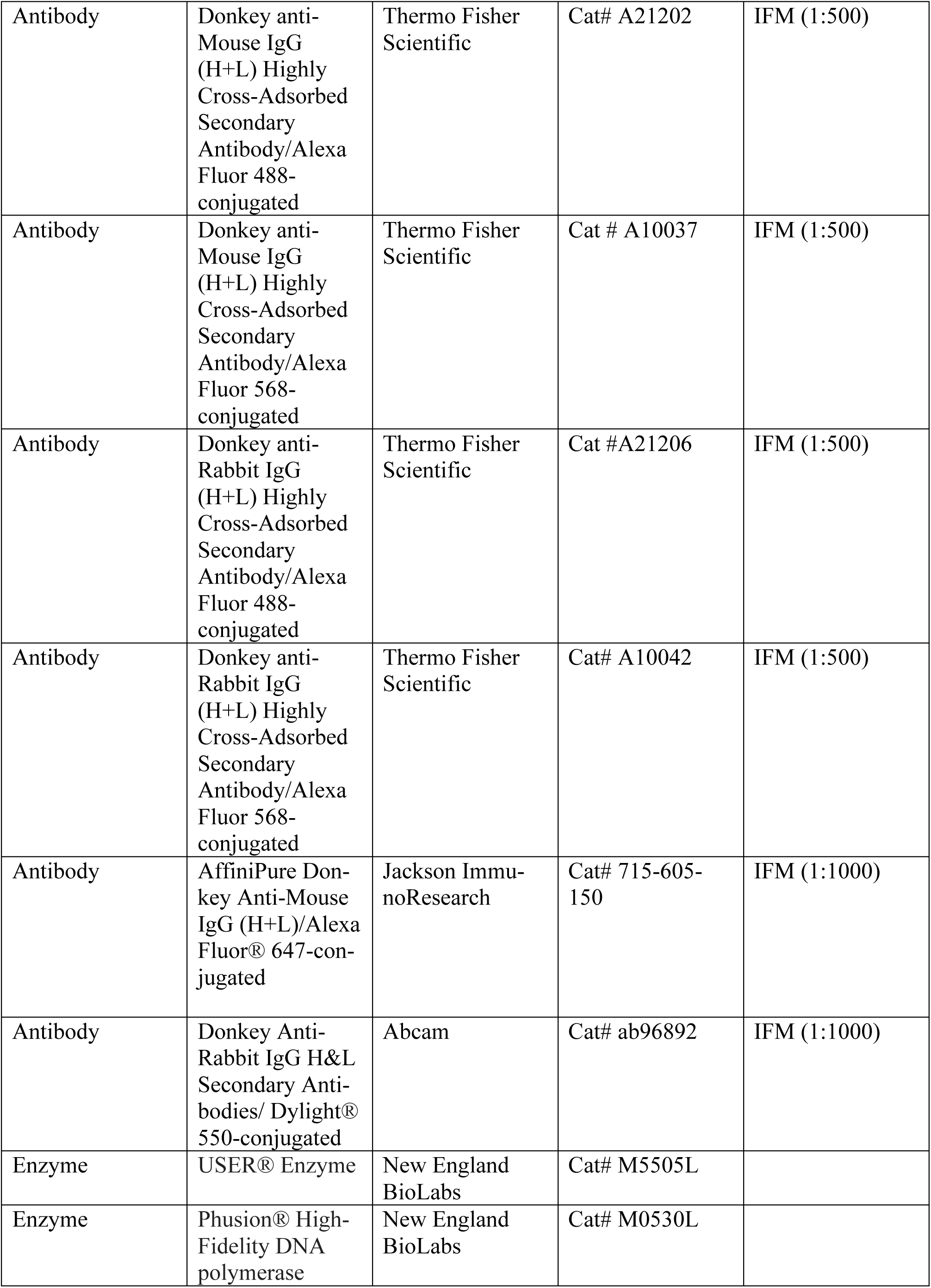
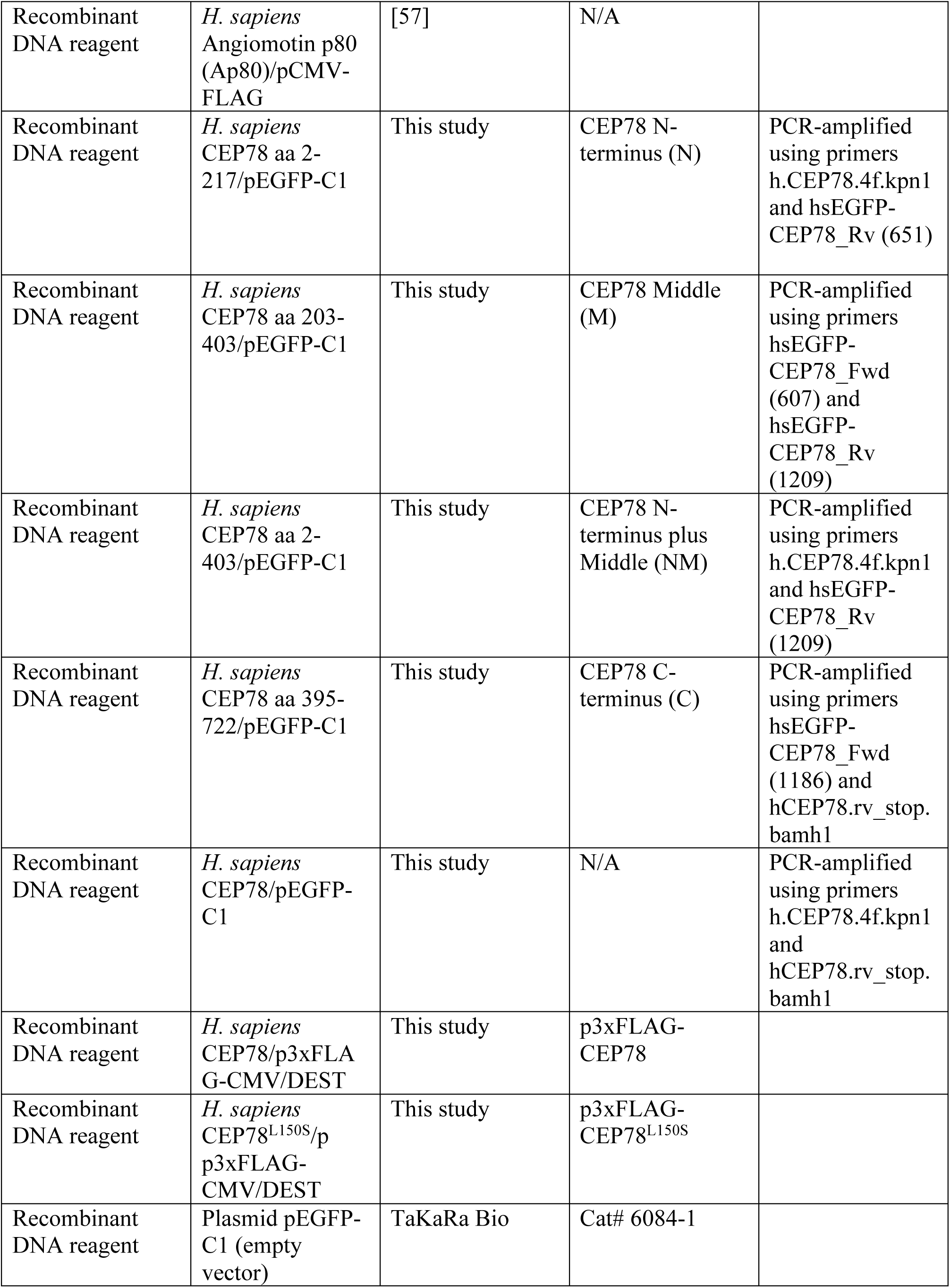
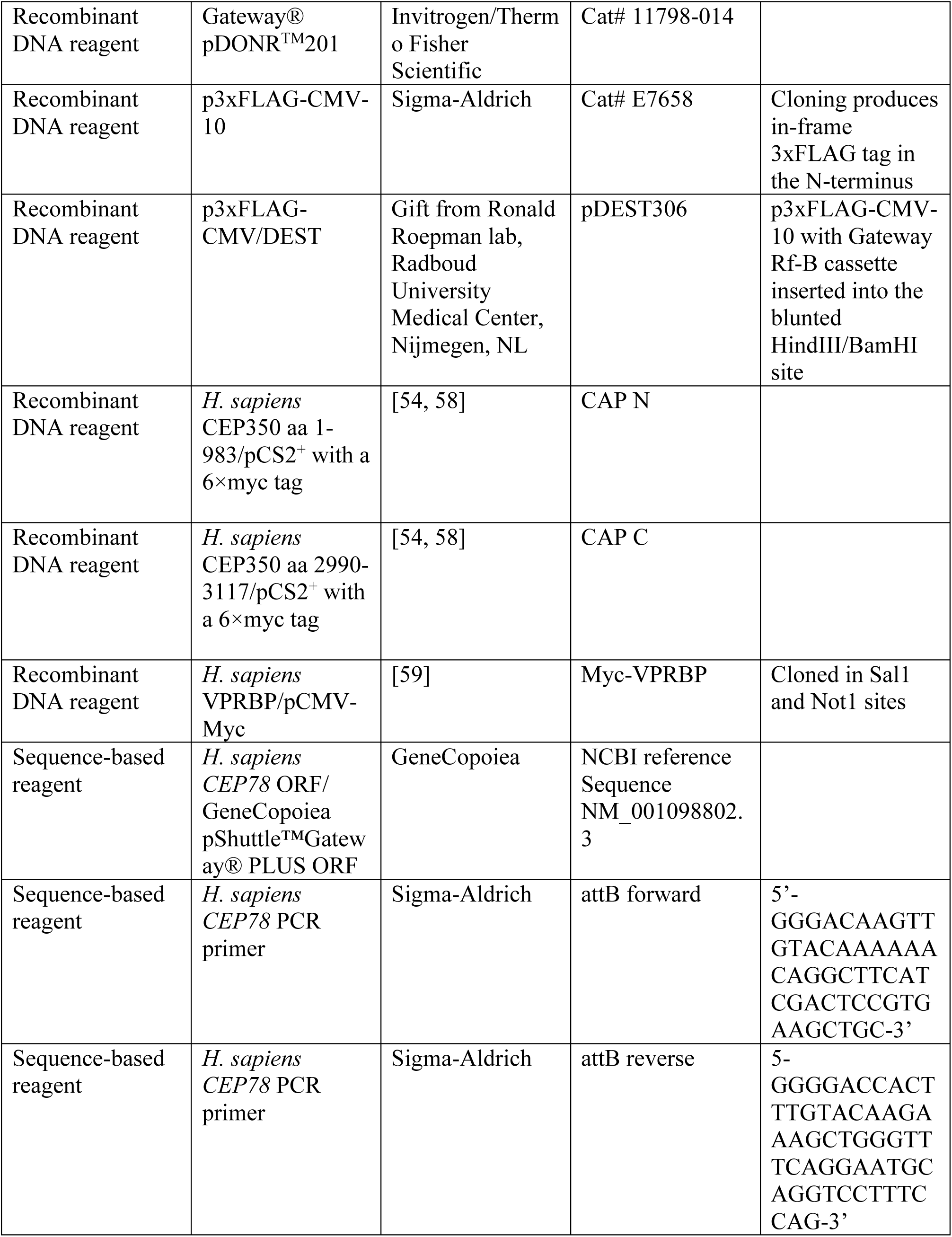
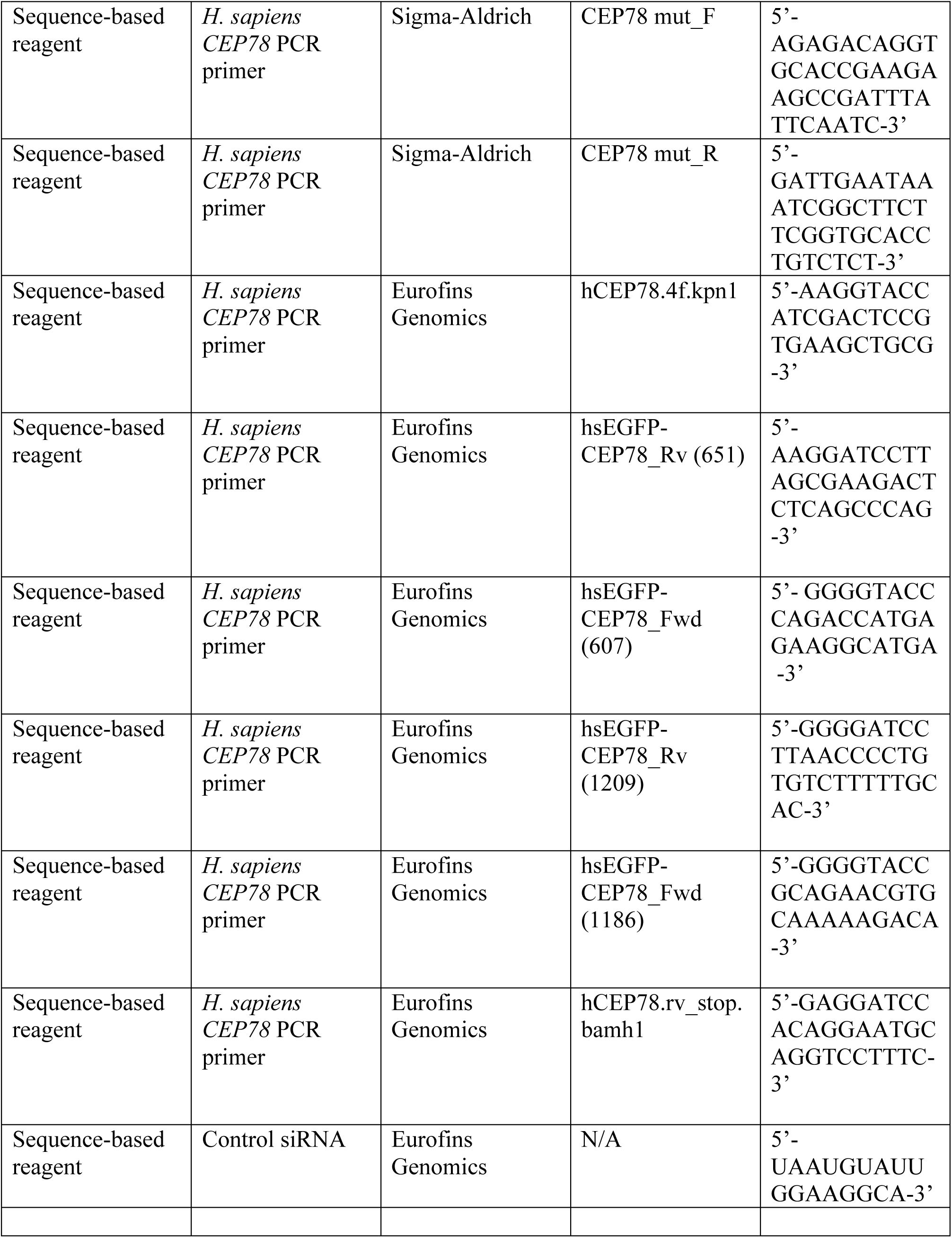
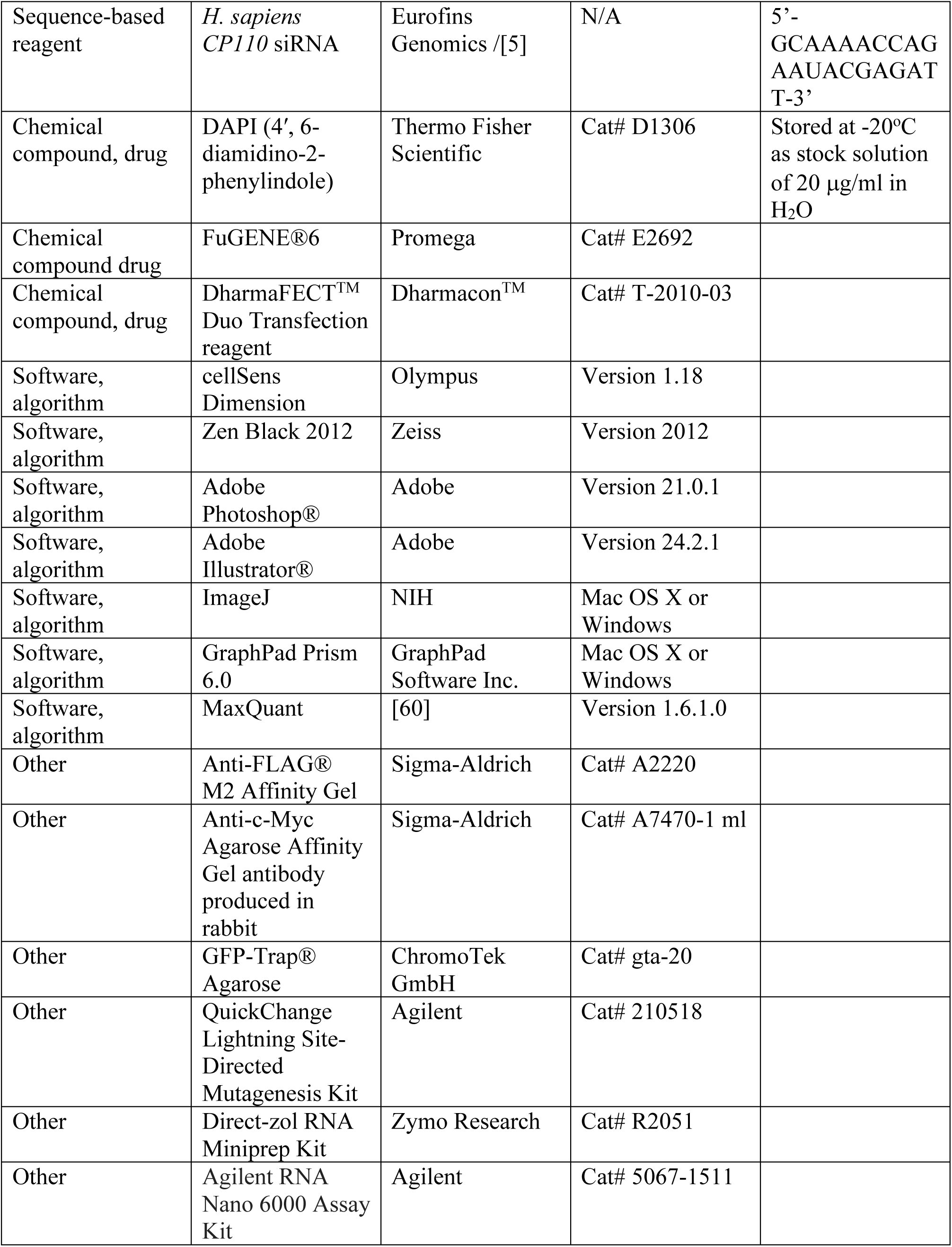
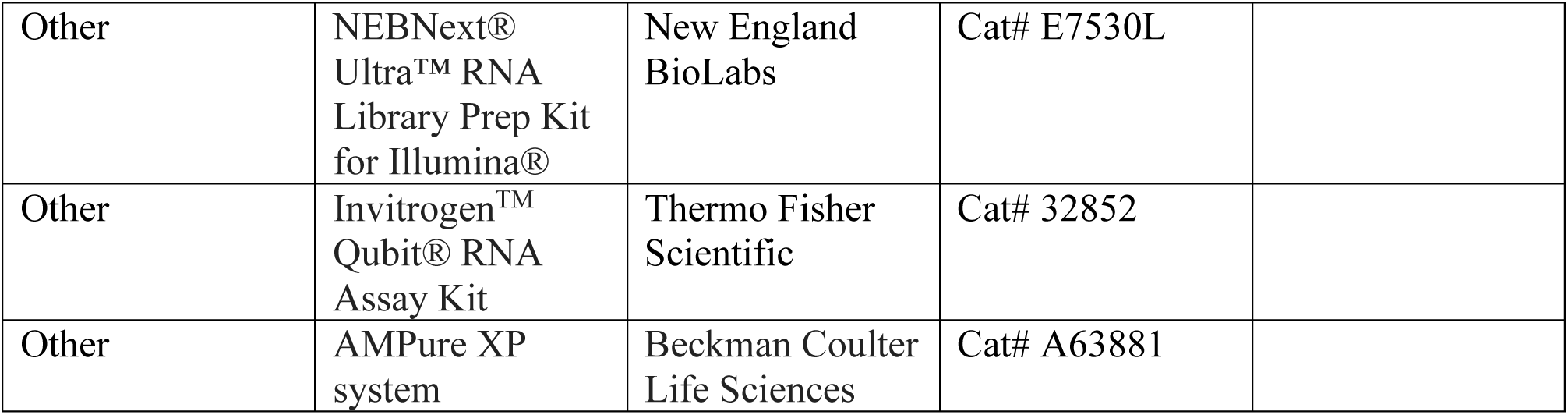

