## Supplemental figures for "CEP78 functions downstream of CEP350 to control biogenesis of primary cilia by negatively regulating CP110 levels"

### Figure 1-figure supplement 1

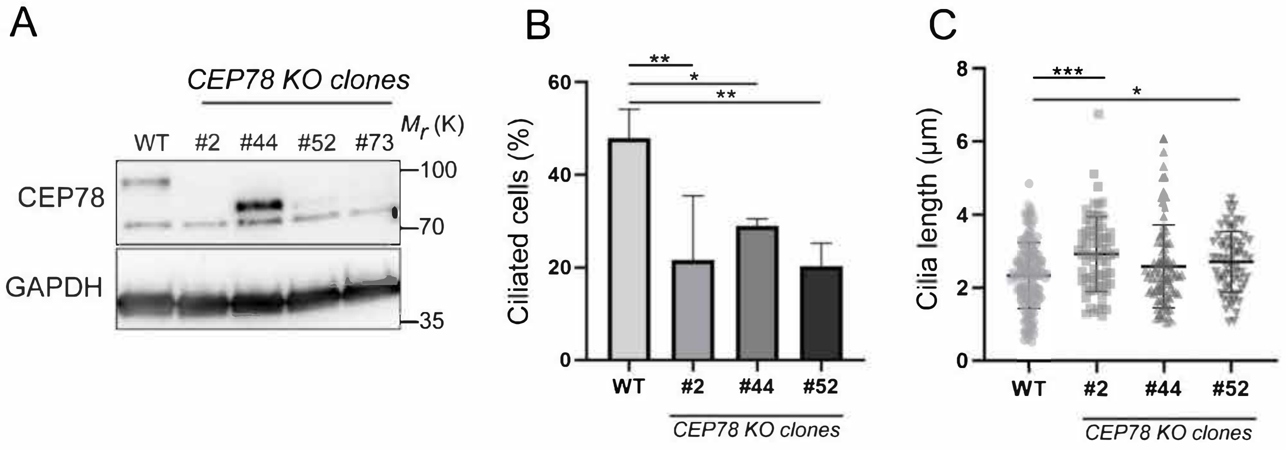

Figure 2-figure supplement 1.

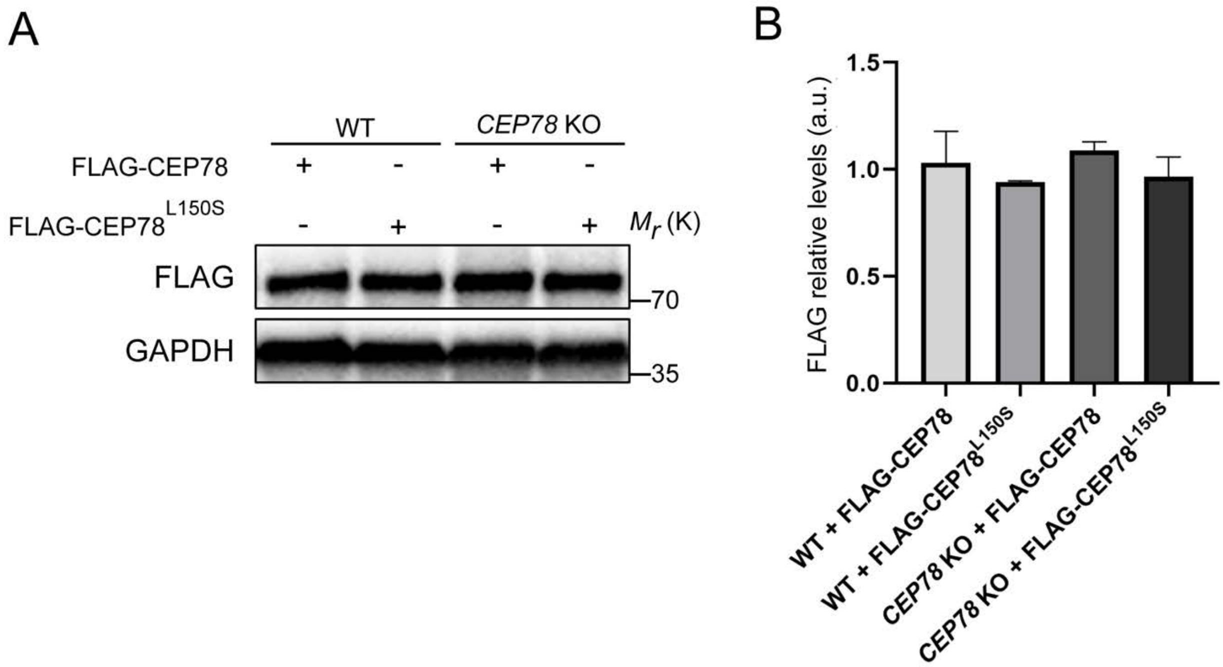

Figure 2-figure supplement 2.

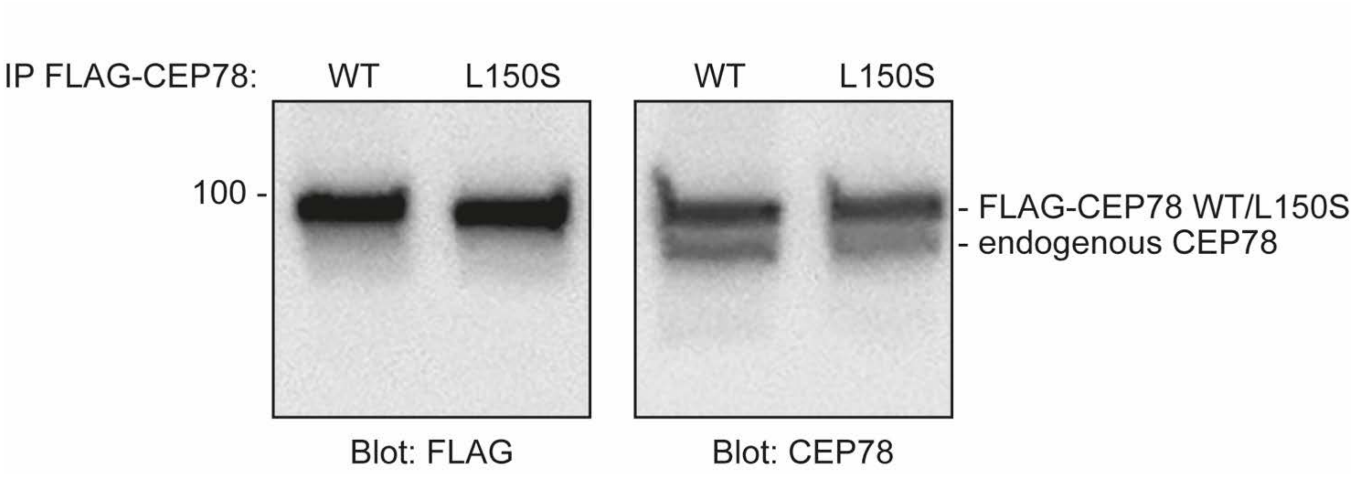

Figure 2-figure supplement 3.

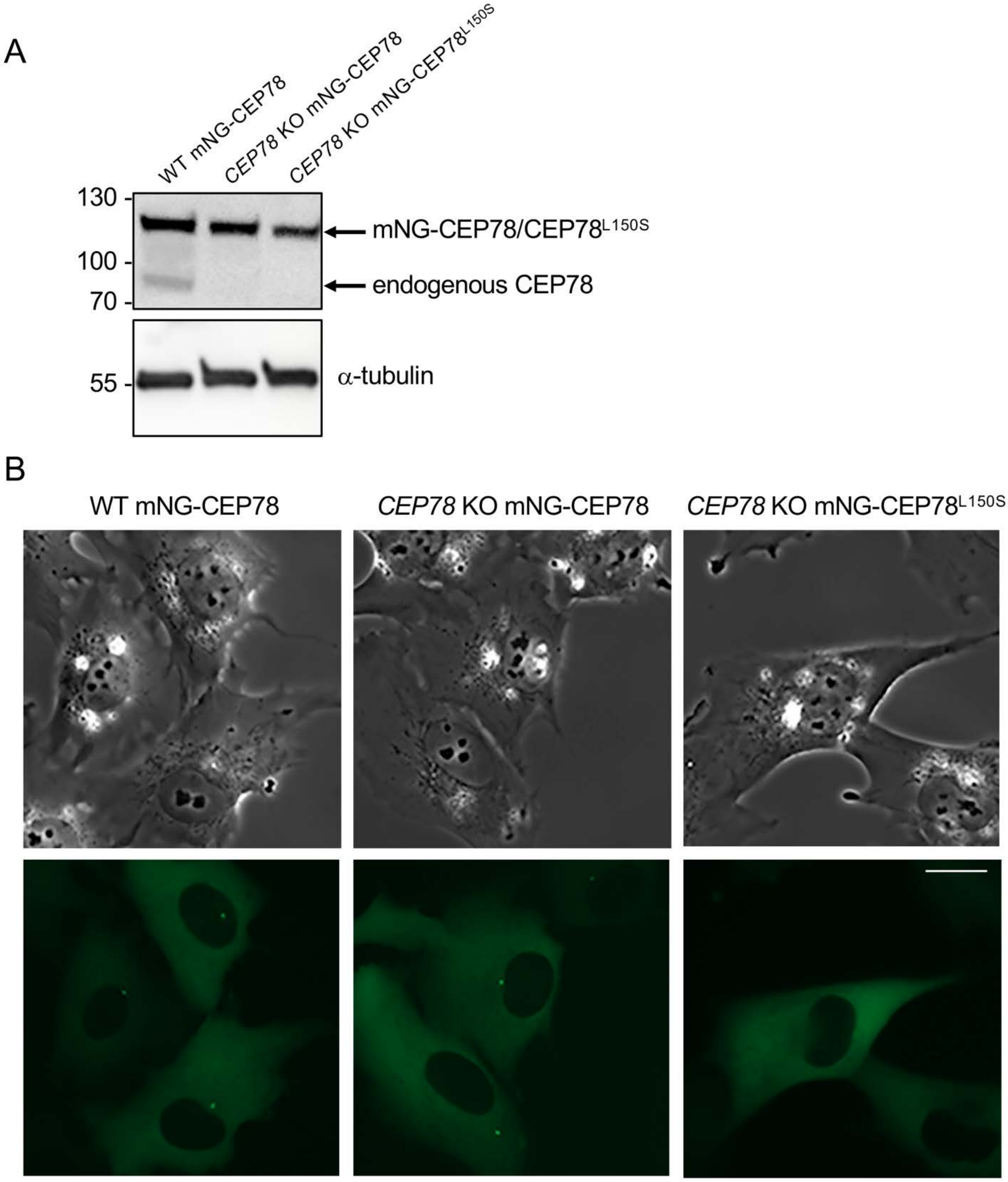

Figure 3-figure supplement 1.

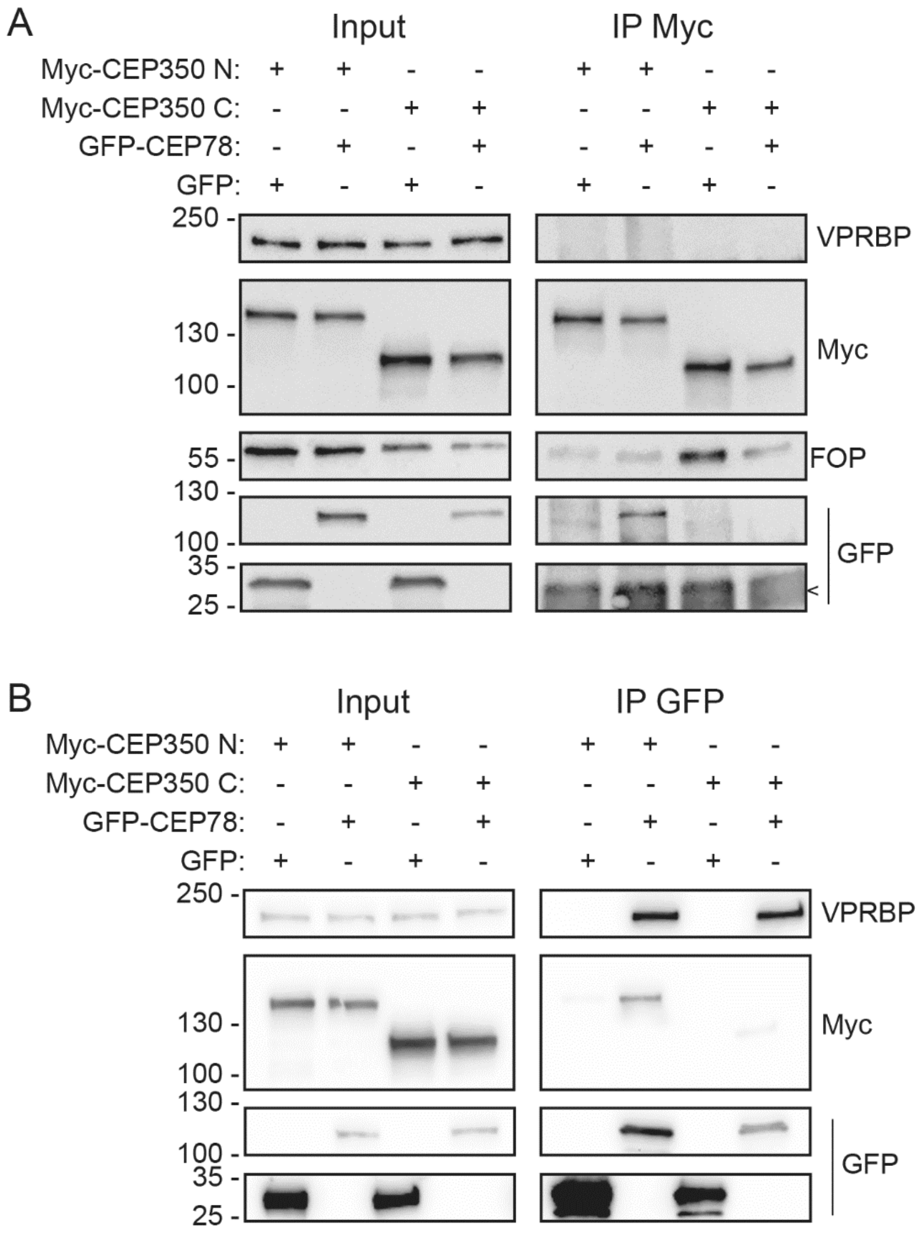

Figure 4-figure supplement 1.

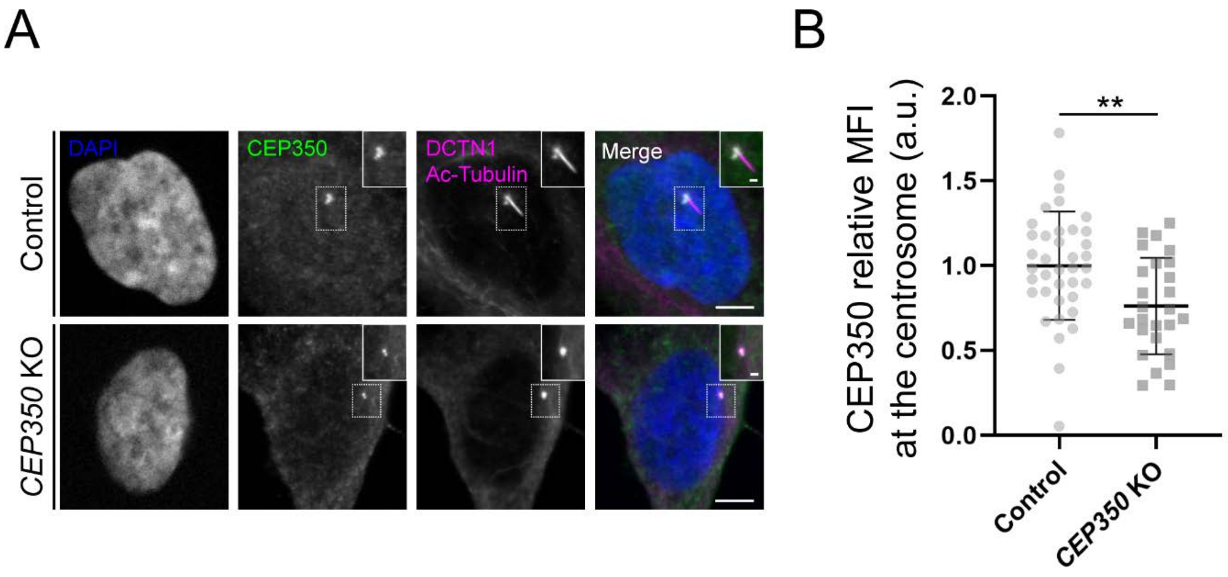

Figure 6-figure supplement 1.

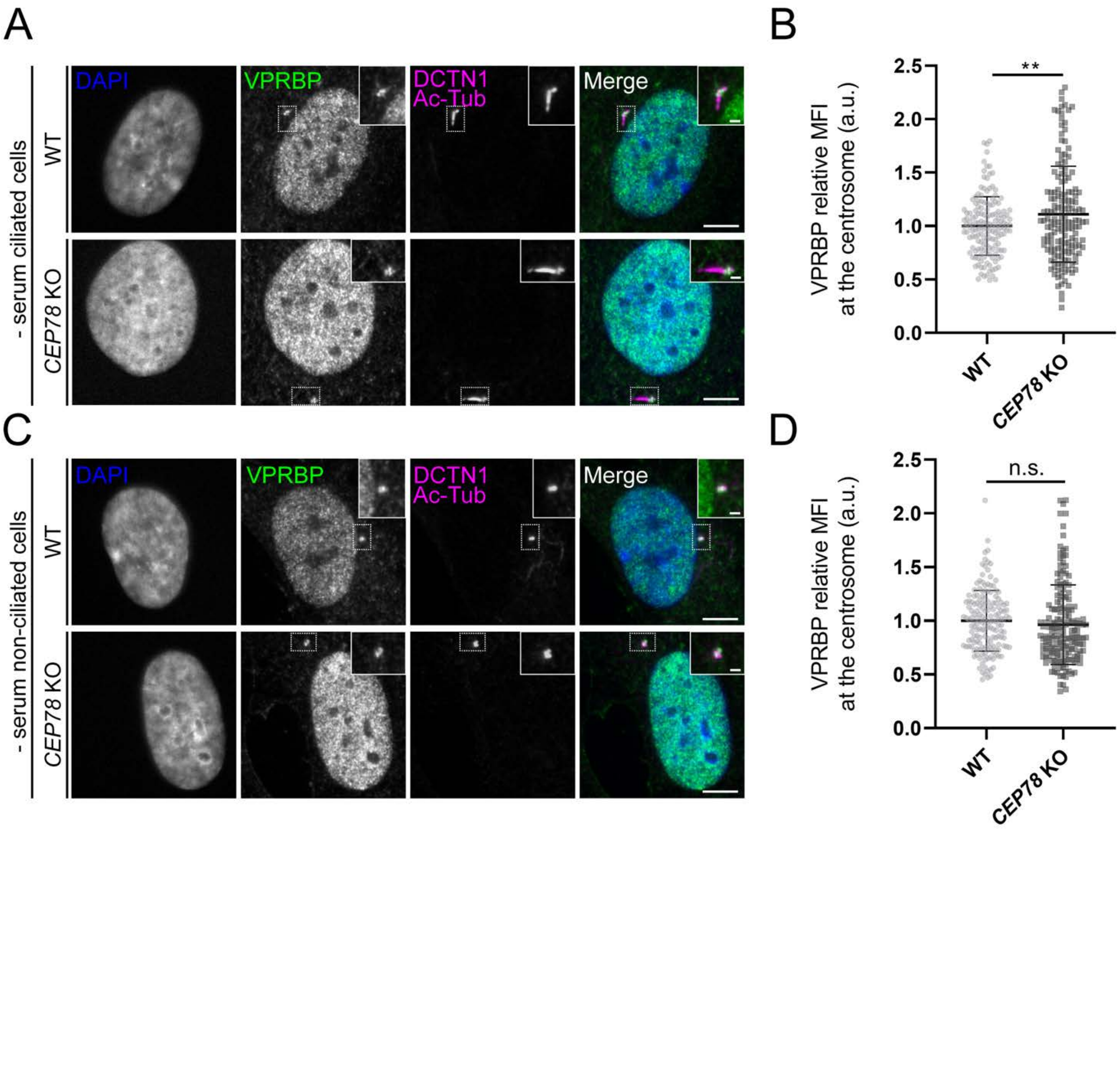

Figure 6-figure supplement 2.

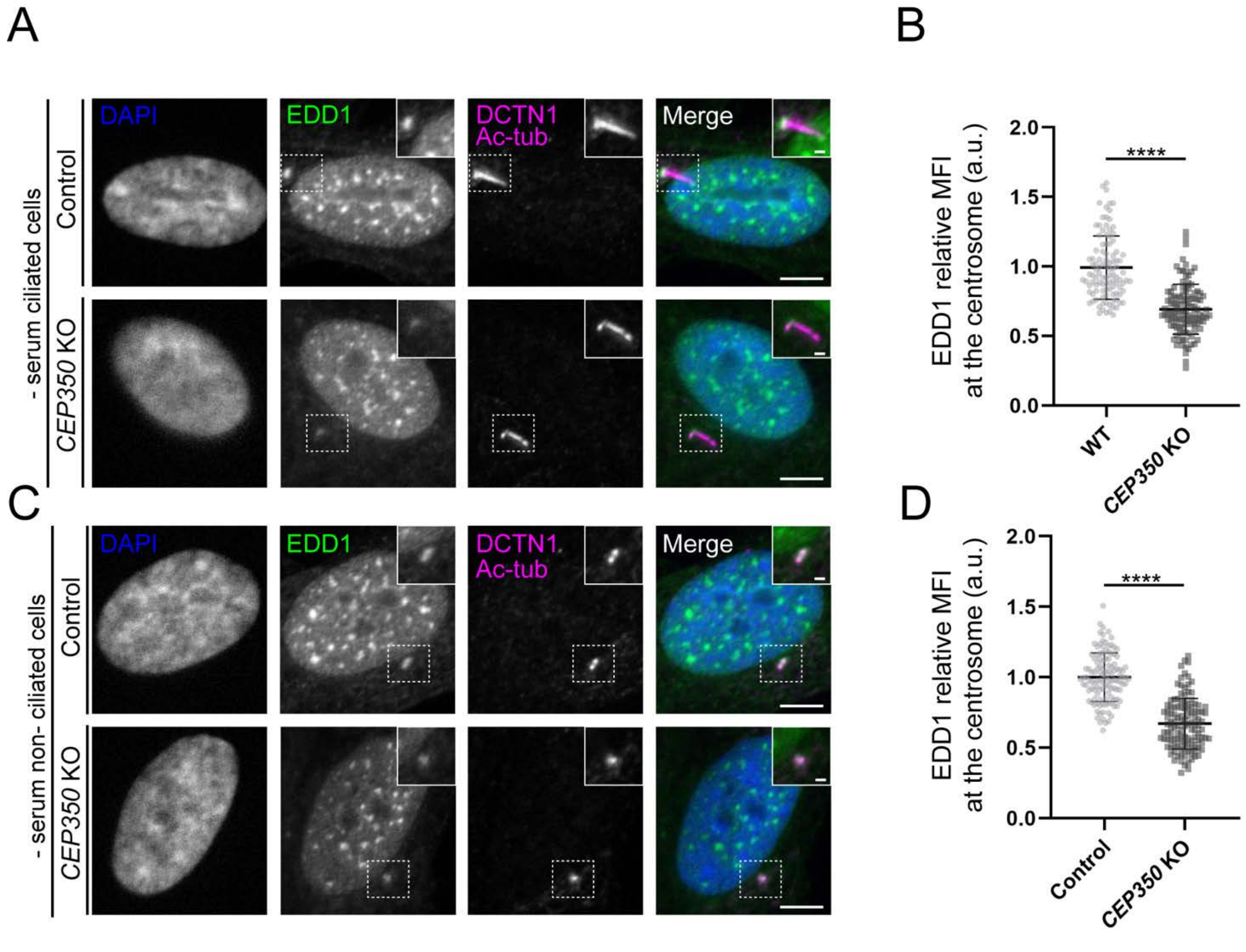

Figure 7-figure supplement 1.

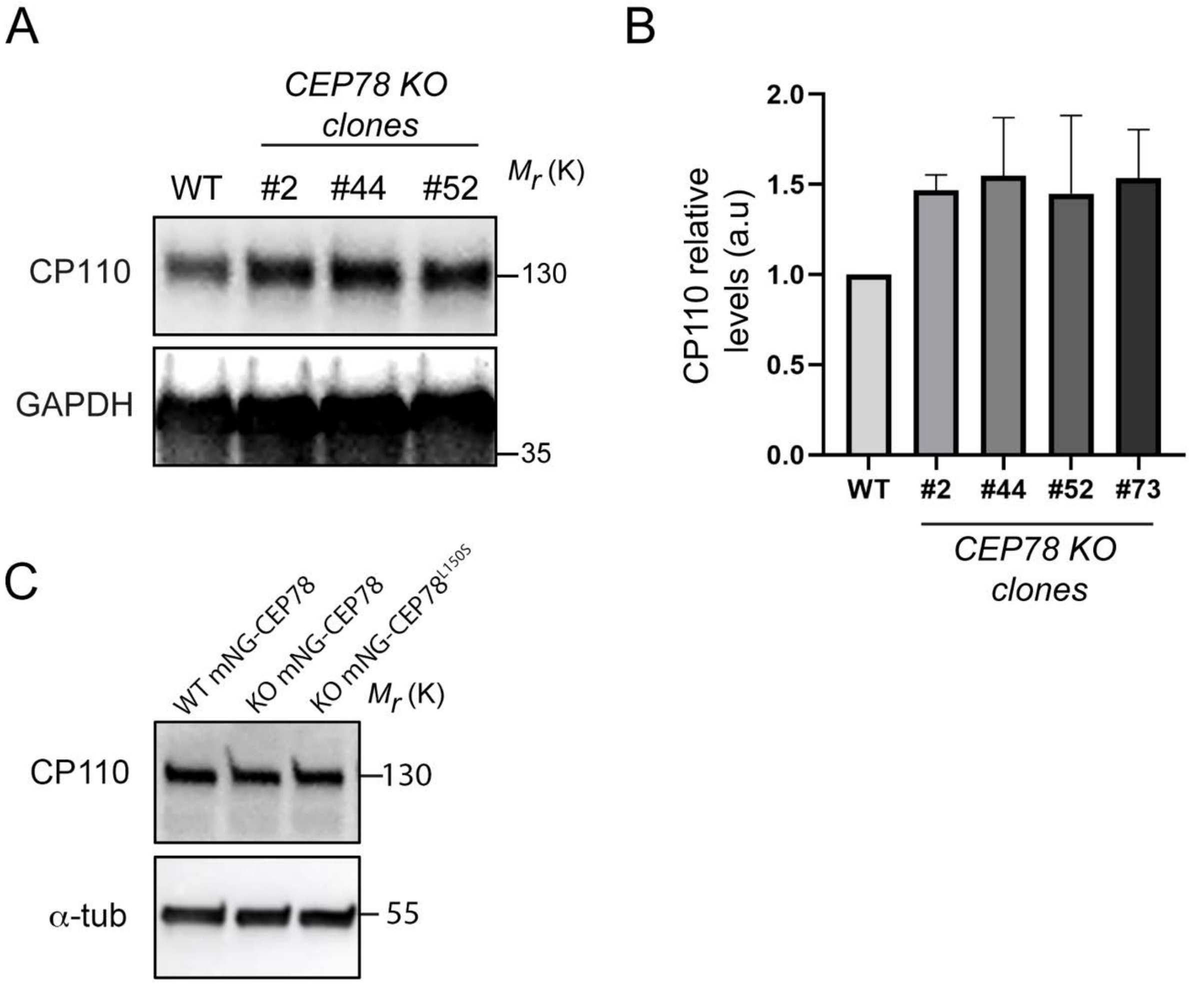

Figure 7-figure supplement 2.

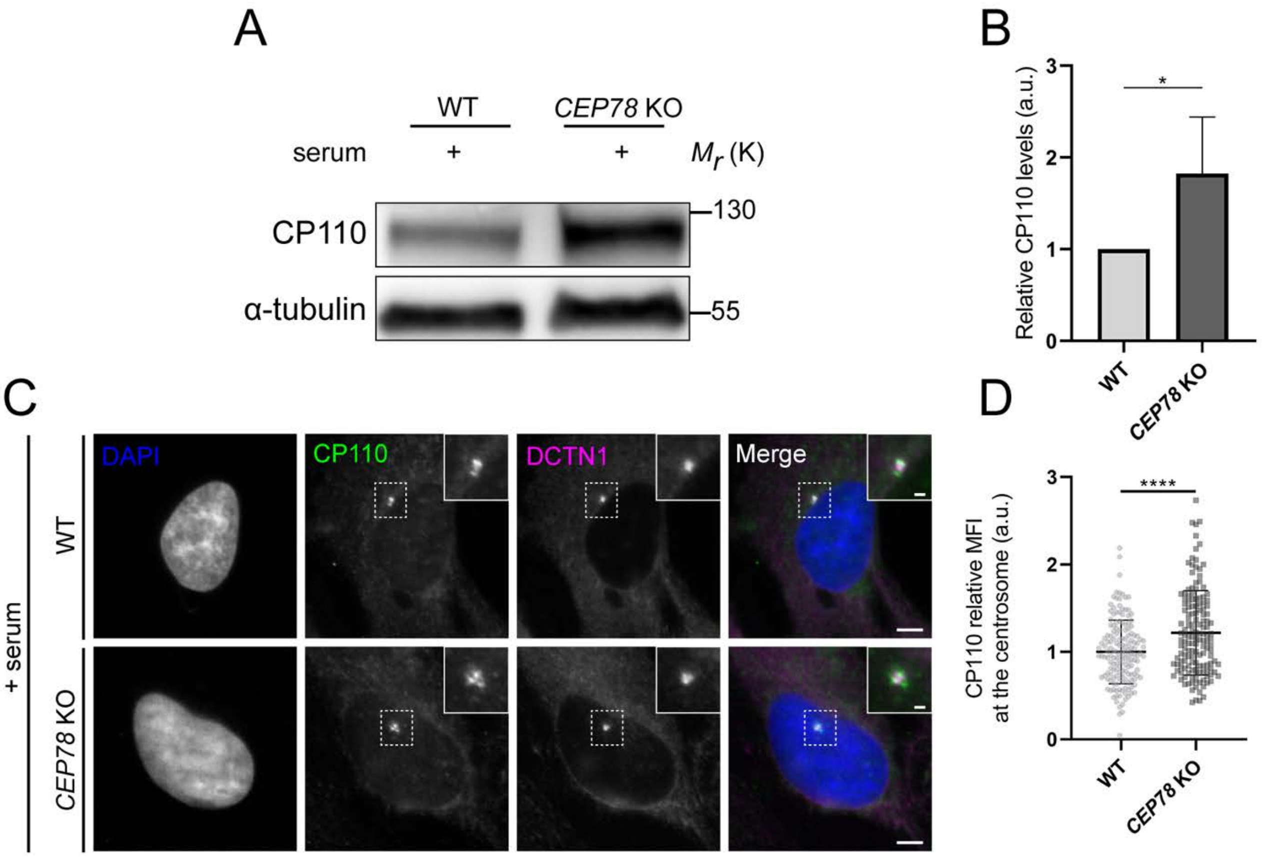

Figure 7-figure supplement 3.

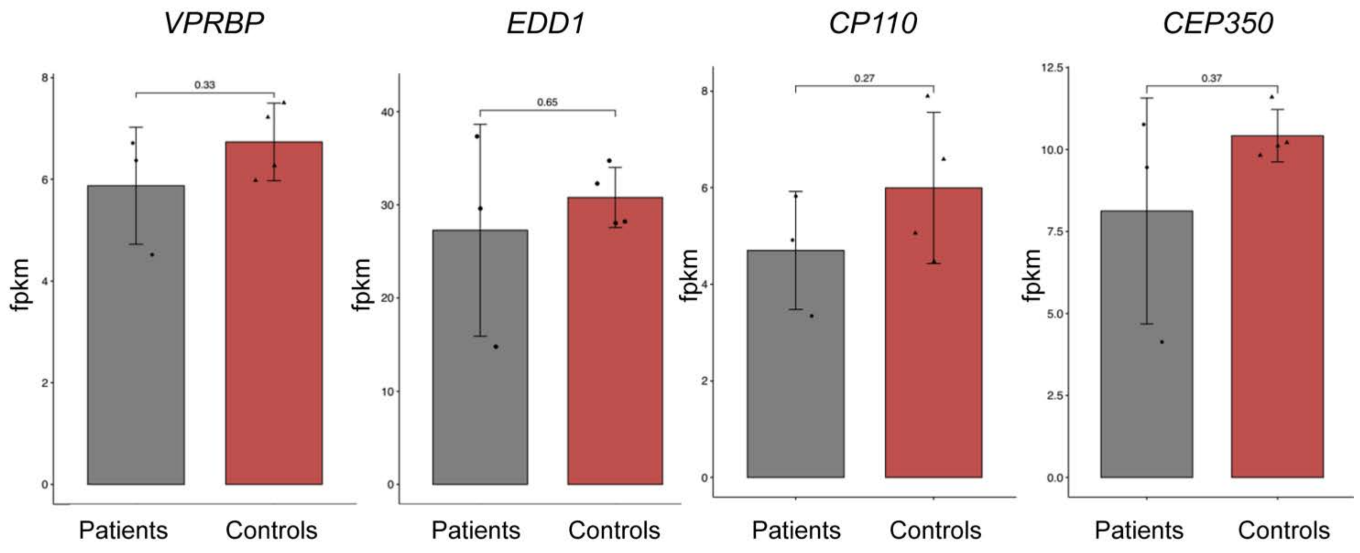

Figure 7-figure supplement 4.

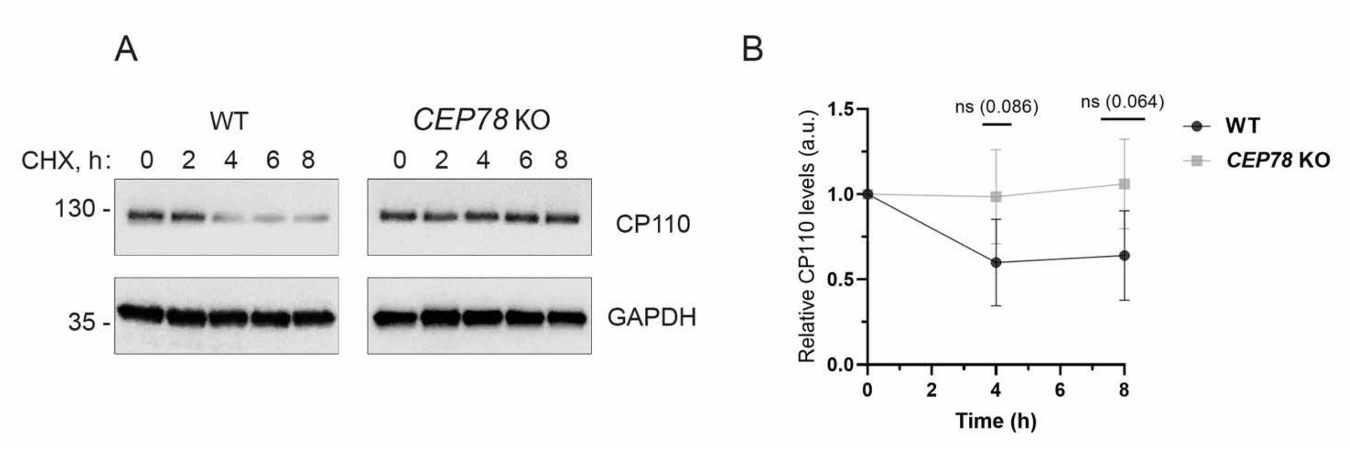

Figure 8-figure supplement 1.

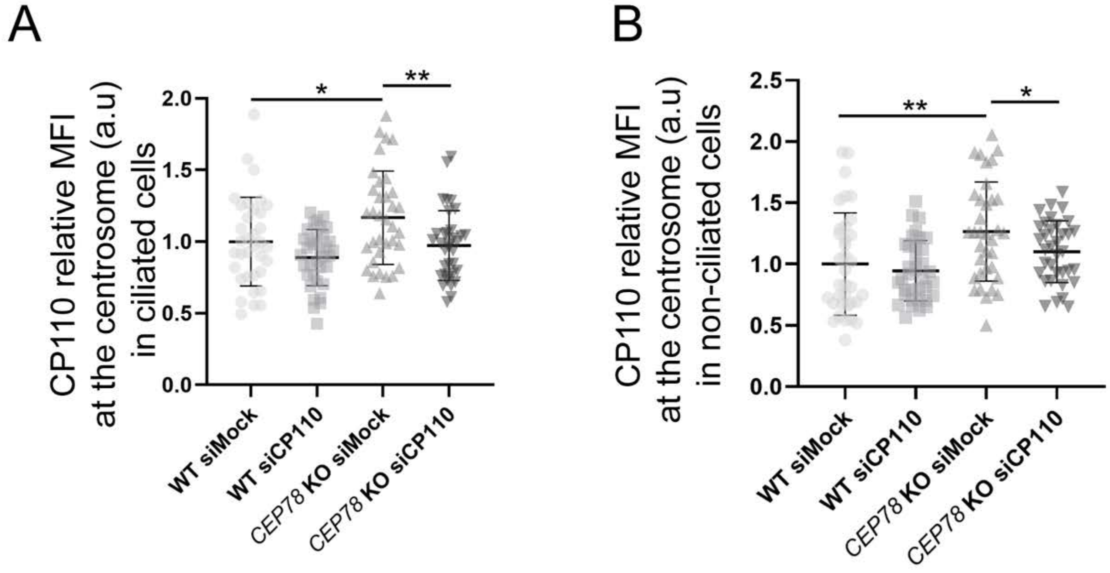
